# Dendritic spine geometry and spine apparatus organization govern the spatiotemporal dynamics of calcium

**DOI:** 10.1101/386367

**Authors:** Miriam Bell, Tom Bartol, Terrence Sejnowski, Padmini Rangamani

## Abstract

Dendritic spines are small subcompartments that protrude from the dendrites of neurons and are important for signaling activity and synaptic communication. These subcompartments have been characterized to have different shapes. While it is known that these shapes are associated with spine function, the specific nature of these shape-function relationships is not well understood. In this work, we systematically investigated the relationship between the shape and size of both the spine head and spine apparatus, a specialized endoplasmic reticulum compartment in the spine head, in modulating rapid calcium dynamics using mathematical modeling. We developed a spatial multi-compartment reaction-diffusion model of calcium dynamics in three dimensions with various flux sources including N-methyl-D-aspartate receptors (NMDAR), voltage sensitive calcium channels (VSCC), and different ion pumps on the plasma membrane. Using this model, we make several important predictions – first, the volume-to-surface area ratio of the spine regulates calcium dynamics, second, membrane fluxes impact calcium dynamics temporally and spatially in a nonlinear fashion, and finally the spine apparatus can act as a physical buffer for calcium by acting as a sink and rescaling the calcium concentration. These predictions set the stage for future experimental investigations of calcium dynamics in dendritic spines.

## Introduction

Dendritic spines, small protein- and actin-rich protrusions located on dendrites of neurons, have emerged as a critical hub for learning, memory, and synaptic plasticity in both short-term and long-term synaptic events [1, 2]. These subcompartments provide valuable surface area for cell-cell interaction at synapses, and compartmentalization of signaling proteins to control and process incoming signals from the presynaptic terminal [3, 4]. Thus, dendritic spines are hotbeds of electrical and chemical activity. Since calcium is the first incoming signal into the postsynaptic terminal, calcium temporal dynamics have been extensively studied experimentally and more recently computationally [4–8]. In particular, calcium acts as a vital second messenger, triggering various signaling cascades that can lead to long term potentiation (LTP), long term depression (LTD), actin cytoskeleton rearrangements, and volume expansion, amongst other events [1, 2, 9].

**Table 1.** Notation used in this study

| Variable/Parameter | Units | Definition/Meaning |
| --- | --- | --- |
| $Ca_{cyto}^{2+}$ | $\mu M$ | Calcium in the cytoplasm |
| $Ca_{ER}^{2+}$ | $\mu M$ | Calcium in the endoplasmic reticulum (spine apparatus) |
| $D$ | $\frac{\mu m^2}{s}$ | Diffusion rate |
| $B_m$ | $\mu M$ | Mobile buffers |
| $B_f$ | $\frac{mol}{\mu m^2 \cdot s}$ | Fixed buffers |
| $K_D$ | $\mu M$ | Dissociation constant |
| $PDE$ | | Partial differential equation |
| $ODE$ | | Ordinary differential equation |
| $J_x$ | $\frac{mol}{\mu m^2 \cdot s}$ | Boundary flux due to x |
| $CBP$ | $\mu M$ | Calcium Binding Proteins (calcium buffers) |
| $V_{cyto}$ | $\mu m^3$ | Volume of the cytoplasm excluding the spine apparatus volume |
| $PM$ | | Plasma membrane |
| $SpApp$ | | Spine apparatus |
| $ECS$ | | Extracellular Space |
| $n$ | $\mu m$ | Scale factor to convert between volume reactions to boundary flux |
| $n_{PM}$ | $\mu m$ | Volume-to-surface area ratio of the cytoplasm to the plasma membrane |
| $n_{SpApp}$ | $\mu m$ | Volume-to-surface area ratio of the SpApp volume to the SpApp surface area |
| AUC | $ions \cdot s$ | Area under the curve |
| EPSP | mV | Excitatory postsynaptic potential |
| BPAP | mV | Back propagating actin potential |
| NMDAR |  | N-methyl-D-aspartate receptor |
| VSCC |  | Voltage sensitive calcium channel |
| PMCA |  | plasma membrane calcium ATPase |
| NCX |  | sodium/calcium exchanger |
| SERCA |  | sarco/endoplasmic reticulum Ca<sup>2+</sup>-ATPase |

Dendritic spine activity has numerous timescales with signaling pathways operating on the millisecond to the hour timescale following spine activation [1, 10, 11]. Calcium dynamics are on the millisecond timescale, since calcium is the second messenger that floods the spine following the release of neurotransmitter from the presynaptic terminal. The temporal dynamics of calcium have provided valuable insight into the signaling dynamics in dendritic spines and it is quite clear that calcium dynamics are influenced by a large number of factors. Multiple studies have connected the electrical activity of the plasma membrane voltage to calcium dynamics of N-methyl-D-aspartate receptors (NMDAR) [12–14]. The electrophysiology of dendritic spines influences many signaling dynamics through voltage-sensitive (or voltage-dependent) ion channels [15] and thus models of these dynamics can be linked to downstream signaling.

Calcium is critical for almost all the reactions in the brain [8] and is believed to accomplish a vast variety of functions through localization [16–18]. One possible way to achieve localization is by restricting the distance between the calcium source (often a channel) and the sink (calcium sensors and buffers). Thus, the localization of calcium can result from the location and mobility of different buffers and sensors [19]. Spatial models of calcium dynamics in dendritic spines that consider such effects have been proposed previously [6, 20, 21]. Spatial-temporal models of calcium dynamics have highlighted the role of calcium-induced cytosolic flow and calcium influx regulation by Ca^2+^-activated K^+^ channels (SK channels) [14, 20]. Computational studies using stochastic models of calcium transients have revealed that the readout of fluorescent probes can alter experimental readouts [6, 10, 22]. In particular, fluorescent probes can sequester calcium, effectively acting as calcium buffers and lowering the perceived calcium concentrations. Therefore, computational studies have already provided some insight into the spatio-temporal dynamics of calcium in both dendritic spines [1, 10, 11] and whole neurons [23]. In this study, we specifically focus on the effect of spine geometry on calcium signals in the postsynaptic spine.

Recent advances in imaging and reconstruction techniques have shed light into the complex surface area of a single spine and the ultrastructure of the spine apparatus [6, 24–26]. Experimental evidence shows that calcium signals remain predominantly localized to single spines [9, 27, 28]. Dendritic spines have a unique set of shapes and recently the size and shape distribution of spines has been linked to their physiological function [29, 30]. Additionally, only about 14% of spines have a specialized endoplasmic reticulum known as the spine apparatus (SpApp) [31, 32], which serves as a calcium store [31, 33]. Indeed, calcium dynamics in dendritic spines are quite complex.

Given the importance of calcium dynamics in dendritic spines and the complexity of spine ultrastructure [24, 34, 35], we sought to use a computational approach to probe the role of spine geometry in modulating the spatio-temporal dynamics of calcium. Specifically, we seek to address the following questions: i) How does the size and shape of both the spine head and spine apparatus affect calcium dynamics? ii) How does the distribution of channels along the synaptic membrane modulate calcium dynamics within the spine? And iii) How do calcium buffers and calcium diffusion rates affect the spatiotemporal dynamics of calcium? To answer these questions, we develop a spatial 3D reaction-diffusion model with multiple compartments. We chose a computational approach because it is not yet possible to manipulate spine size, shape, or spine apparatus location with precise control experimentally. However, the insights obtained from computational approaches can lay the groundwork for generating experimentally testable predictions [36].

### Model assumptions

In order to interrogate the spatiotemporal dynamics of calcium in dendritic spines, we developed a reaction-diffusion model that accounts for the fluxes through the different sources and sinks shown in Figure 1. We briefly discuss the assumptions made in developing this model and outline the key equations below.

**Fig 1.**
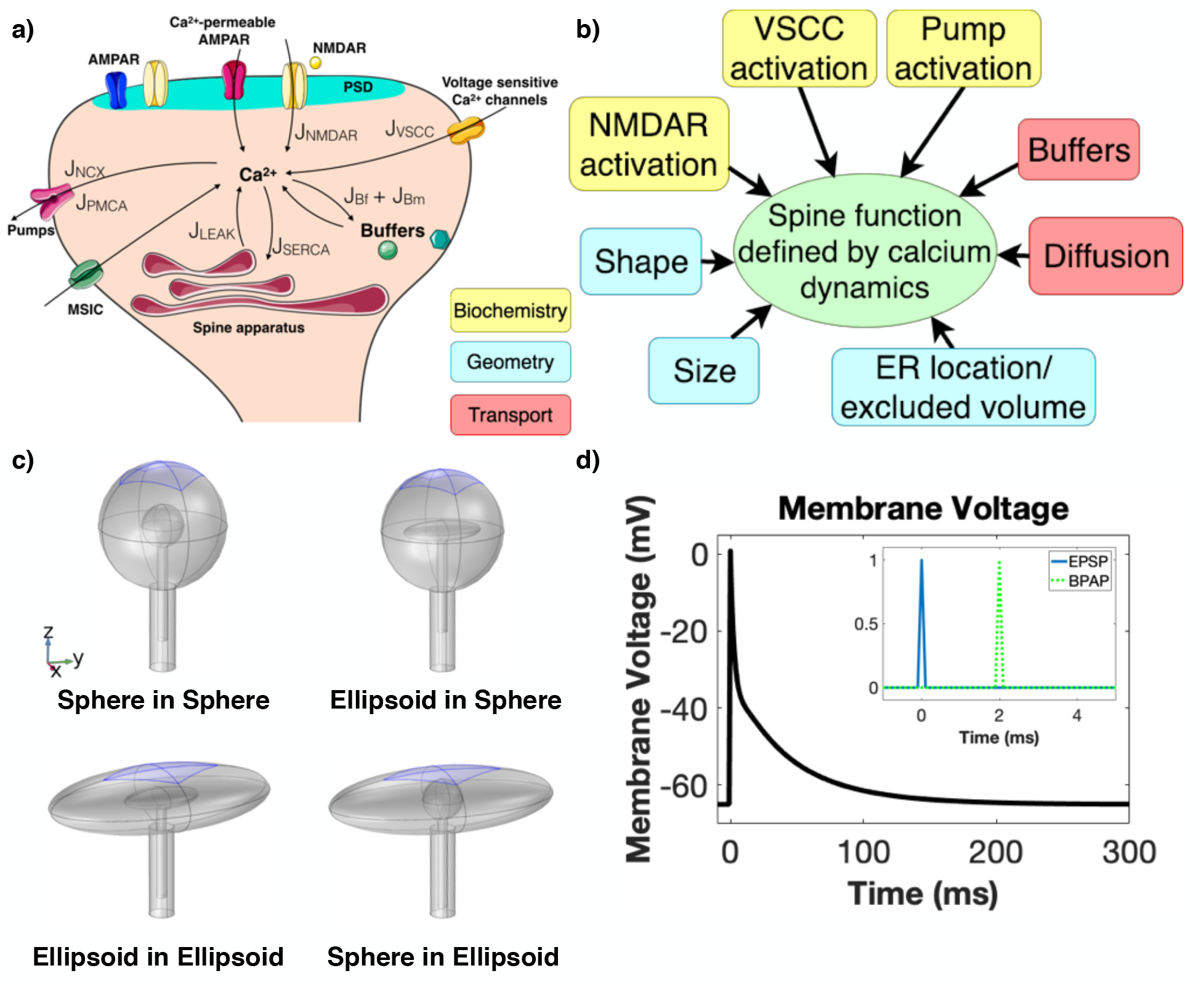
Physical and chemical determinants of calcium influx in dendritic spines. a) Spatiotemporal dynamics of calcium in dendritic spines depend on multiple sources and sinks on both the spine membrane and the spine apparatus membrane. These include receptors (NMDAR), channels (VSCC), and pumps (plasma membrane calcium ATPase (PMCA), sodium/calcium exchanger (NCX)). Calcium buffers are present in both the cytoplasm and on the plasma membrane. b) A partial list of factors that can influence these dynamics include biochemical components (shown in panel a), geometry, and protein transport components, which are effectively coupled through transport phenomena. In this study, we focus on the effects of spine and spine apparatus size, spine and spine apparatus shape, flux through NMDAR and VSCC distribution on calcium spatiotemporal dynamics, and buffers. c) Four different combinations of spine head and spine apparatus geometries are used as model geometries – spherical head with spherical apparatus, spherical head with ellipsoidal apparatus, ellipsoidal head with ellipsoidal apparatus, and ellipsoidal head with spherical apparatus – to study how spine geometry affects calcium dynamics. The coordinate axes correspond to 100 nm in the different geometries. The blue shaded regions denote the PSD for each geometry. d) In our model, depolarization of the membrane is triggered by an excitatory postsynaptic potential (EPSP) followed closely by a back propagating action potential (BPAP) to create a maximal depolarization according to spike time dependent plasticity (STDP). This membrane voltage acts as the input to our model. Inset: Timing of the EPSP and the BPAP. We model the maximum possible membrane depolarization based on STDP with the EPSP arriving 2 ms before the BPAP.

1. **Time scale**: We model a single dendritic spine of a rat hippocampal area CA1 pyramidal neuron as an isolated system because we focus on the 10-100 ms timescale and the ability of the spine neck to act as a diffusion barrier for calcium [7, 16, 37, 38]. As a result, we do not consider calcium dynamics due to the mitochondria. This assumption is valid in our model because even though mitochondria are known to act as calcium stores, their dynamics are on a longer timescale (10 - 100 s) [39] and mitochondria are located in the dendrite outside of the dendritic spine. Although, it is well-known that the spine apparatus acts as a source for calcium, Ca^2+^ release from the spine apparatus occurs at longer timescales because of IP_3_ receptors, Ryanodine receptors, and calcium-induced calcium release [31]. It should also be noted that not all neurons have RyR and IP_3_R on the spine apparatus [40]. Therefore, for the purpose and timescale of this study, we do not focus on calcium-induced calcium release (CICR), and therefore do not include RyR or IP_3_R dynamics in this study.
2. **Membrane voltage stimulus**: We model membrane voltage as an algebraic equation based on the summation of an EPSP and a BPAP applied to the entire plasma membrane [12, 13, 41]. The EPSP arrives 2 ms before the BPAP to stimulate the maximum possible membrane depolarization (Figure 1d) [42]. This stimulus triggers the influx of calcium into the spine head.
3. **Spine head**: The average spine head volume is approximately 0.03 *µm*^3^ [6,34], but a large variation has been observed physiologically. The commonly observed shapes of dendritic spines include filopodia-like, stubby, short, and mushroom-shaped spines [2, 30]. In this work, we consider two idealized geometries – a spherical spine head to represent a younger spine, and an ellipsoidal spine head to represent a more mature mushroom spine [34]. The postsynaptic density (PSD) is modeled as a section of the membrane at the top of the spine head (Figure 1c). In simulations where the spine size is varied, we assume that the PSD changes surface area approximately proportionally to the spine head volume [43].
4. **Spine apparatus**: Spine apparatuses are found primarily in larger mushroom spines [34], which hints at their role in potential regulation of sustained spine volume [44]. We assume that the spine apparatus acts as a calcium sink within the timescale of interest [45]. Another assumption is that the spine apparatus has the same shape as the spine head, a simplification of the more complicated and intricate spine apparatus geometry [34].
5. **Plasma membrane fluxes**: To maintain our focus on the short-time scale events associated with calcium dynamics, we include the following plasma membrane (PM) sources and sinks of calcium – voltage sensitive calcium channels (VSCC), N-methyl-D-aspartate receptors (NMDAR), plasma membrane Ca^2+^-ATPase (PMCA) pumps, and sodium-calcium exchanger (NCX) pumps. NMDAR are localized on the postsynaptic membrane adjacent to the postsynaptic density (PSD), designated at the top of the spine head. VSCC, PMCA, and NCX pumps are uniformly distributed along the plasma membrane, including at the base of the spine neck. Therefore, we model the dendritic spine as an isolated system with the spine neck base modeled in the same manner as the rest of the PM, rather than explicitly modeling the base with an outward flux into the dendrite (see assumption 6).
6. **Boundary condition at the base of the neck**: We model the spine as an isolated system [7,16,27,46]. Therefore, for most of our analyses, we use the same boundary conditions at the base of the spine neck as the rest of the PM. In the supplemental material (SOM), we relax this assumption and test different boundary conditions including a clamped calcium concentration at the base of the neck and an explicit effect of a dendritic shaft attached to the spine neck (Figures S4 and S5).
7. **Compartmental specific calcium ion concentration**: We explicitly model calcium in the cytoplasm and in the spine apparatus. We assume that the calcium concentration in the extracellular space (ECS) is large enough (2 mM) [6, 17] that the calcium influx into the spine has an insignificant effect on ECS calcium concentration. Therefore, ECS calcium concentration is assumed constant. We only solve the volumetric reactions in the cytoplasm. The ER calcium concentration is assumed to only affect the fluxes on the ER membrane.
8. **Calcium binding proteins – Buffers**: There are numerous calcium binding proteins (CBPs) present in the cytoplasm that act rapidly on any free calcium in the spine head [6, 10, 16, 27, 47]. These CBPs are modeled as both mobile and fixed buffers in our system. Mobile buffers are modeled as volume components in the cytoplasm [19, 48], and they are modeled as a diffusive species with mass-action kinetics in the cytoplasm. We assume that the mobile buffers have a buffering capacity, *κ*, of 20, where 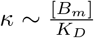 [27, 49] (*K*_*D*_ is the dissociation constant, *B*_*m*_ is the mobile buffer concentration). Dendritic spines also have fixed or immobile buffers but the molecular identity of fixed buffers remains more elusive. Studies suggest that they are primarily membrane-bound components [49]; therefore, we model fixed buffers as immobile species localized to the plasma membrane. As a result, the interactions of calcium with these fixed buffers are treated as flux boundary conditions for the surface reactions with mass-action kinetics. We assume fixed buffers have a dissociation constant of *K*_*D*_ = 2*µM* [6] and have a concentration of 78.7*µM* [6]. These values are converted to a membrane density by multiplying by the spine volume over PM surface area (*n_PMr_*) (see SOM Section S1.1 for more details.).
9. **Spine apparatus fluxes**: In this model, the spine apparatus acts as a Ca^2+^ sink in the 10-100 ms timescale [31, 50, 51]. The implications of this assumption are discussed later in the manuscript. We assume that sarco/endoplasmic reticulum Ca2+-ATPase (SERCA) pumps are located uniformly on the spine apparatus membrane. SERCA pumps have been observed to buffer a large percentage of calcium within the spine, so we include SERCA pumps with a relatively large influx [22, 42]. We also include a small leak current from the spine apparatus to the cytoplasm and set this leak current to offset pump dynamics at basal calcium concentrations of 100 nM [6, 52].

Based on these assumptions, we constructed a 3-dimensional spatial model of Ca^2+^ dynamics in dendritic spines. Our control geometry is a medium-sized spine with volume of ~0.06 *µm*^3^ including the spine head and neck, with a spine apparatus of volume ~0.003 *µm*^3^. We use a spherical spine with spherical spine apparatus and ellipsoidal spine with ellipsoidal spine apparatus as our two control spines of interest. Most results are shown as a 2D cross section for ease of interpretation, see SOM Figure S7 for examples of the full 3D solutions.

### Spatial model of Ca^2+^ influx

The spatiotemporal dynamics of calcium are determined by the combination of dynamics within the spine volume and boundary conditions at the plasma membrane and spine apparatus. Calcium dynamics in the spine volume are represented by a single reaction-diffusion equation

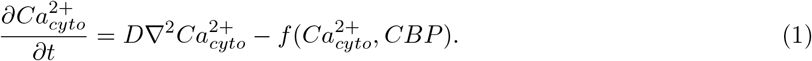

Here, *D* is the diffusion coefficient of calcium and 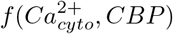 is a function that represents the reaction between cytosolic calcium and mobile buffers in the cytoplasm; ∇^2^ is the Laplacian operator in 3D. The stimulus to the system is the depolarization of the membrane based on an ESPS and BPAP separated by 2 ms, Figure 1d. The boundary conditions at the PM and the spine apparatus are given by time-dependent fluxes that represent the kinetics of different channels, pumps, and fixed buffers.

### Boundary conditions at the PM

We model the calcium influx through activated NMDARs in response to glutamate release in the synaptic cleft and the calcium influx through VSCCs in response to membrane depolarization [6, 12, 13]. We should note that the majority of existing models for NMDAR and VSCC calcium influx assume well-mixed conditions. In this model, these species are restricted to the PM, which is the boundary of the geometry. This results in a time-dependent flux at the PM. Both the NMDAR and VSCC-mediated calcium influx depend on the membrane voltage (see Figure 1d); we prescribe this voltage as a set of biexponentials to capture a back propagating action potential (BPAP) and excitatory post-synaptic potential (EPSP) based on [12, 13]. On the PM, we also include PMCA and NCX that are activated in response to a change in cytosolic calcium concentration [53, 54]. We also localize fixed calcium binding proteins (fixed buffers) to the plasma membrane. Therefore, the binding of cytosolic calcium to fixed buffers (*B*_*f*_) is modeled as a membrane flux, 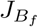. The flux boundary condition at the PM is then the sum of all these fluxes and is given by

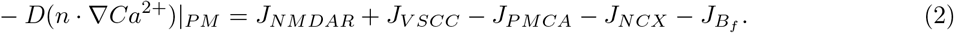

The functions that define the flux terms in Equation (2) are given in Table S1.

### Boundary condition at the spine apparatus membrane

In the cases where we included a spine apparatus, we included SERCA pumps and a leak term along the spine apparatus membrane that are functions of the cytosolic calcium concentration. The boundary condition for the flux across the spine apparatus membrane is given by

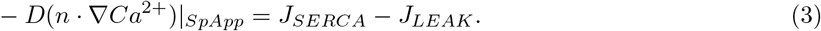

The functions that define the flux term in Equation (3) are given in Table S1.

To briefly summarize, these governing equations are simply the balance equations that keep track of the spatiotemporal dynamics of cytosolic calcium due to calcium diffusion and mobile buffers (Equation (1)), influx and efflux through the PM (Equation (2)), and influx and efflux through the spine apparatus membrane (Equation (3)). The coupled nature of this system of equations and time-dependent fluxes limits the possibility of obtaining analytical solutions even for simple geometries [55]. Therefore, we use computational methods to solve these equations.

### Parametric sensitivity analyses

Given the vast number of parameters in this model, we constrain the parameters in our model as follows: parameter values are chosen from experimental observations or existing computational models, or to match overall experimental and computational observations with respect to pump or channel dynamics (section S1). Overall, we predict a high calcium concentration and relatively fast decay dynamics in accordance with [22, 27, 42]. See Figure S2 for a comparison of temporal dynamics to existing literature. We conducted a kinetic parameter sensitivity analysis for our model using COPASI (S2). The sensitivity analysis was performed in COPASI by converting our spatial model into a compartmental ODE system. This conversion involves transforming boundary flux equations into volumetric reaction rate through the lengthscale factor ‘n’, volume-to-surface area ratio.

### Geometries used in the model

We modeled the dendritic spines using idealized geometries of spheres and ellipsoids; dendritic spines consist of a spine head attached to a neck, with a similarly structured spine apparatus within the spine, see Figure 1c for the different model geometries used in this study. These geometries were inspired by the reconstructions in [6, 41, 56] and were idealized for ease of computation. For the variations of the base of the neck, we also include a condition with an explicit dendrite modeled as a cylinder attached to the spine neck, Figures S4 and S5. The geometric parameters, including volume and surface area, are given in Table S8.

### Numerical Methods

Simulations for calcium dynamics were conducted using commercial finite-element software (COMSOL Multiphysics 5.4). Specifically, the general form and boundary partial differential equations (PDEs) interface were used and time-dependent flux boundary conditions were prescribed. A user-defined tetrahedral mesh with a maximum and minimum element size of 0.0574 *µ*m and 0.00717 *µ*m, respectively, was used. Due to the time-dependent, non-linear boundary conditions used in this model, we also prescribed 4 boundary layers (prism mesh elements) on all membranes in COMSOL. A time-dependent solver was used to solve the system, specifically a MUMPS solver with backward differentiation formula (BDF) time stepping method with a free time stepper. Results were exported to MATLAB for further analysis. All COMSOL files will be posted on the Rangamani Lab website for public dissemination after the manuscript is published.

## Results

Using the model developed above (Equations (1) to (3)), we investigated how different geometric factors of the spine head and the spine apparatus affect calcium dynamics. Because of the coupling between the volume dynamics (Equation (1)) and the fluxes on the membranes due to biochemical components (Equations (2) and (3)), the effect of spine geometry on calcium dynamics is quite complex. In order to parse the coupled effects, we have categorized and organized our simulations as follows and discuss each case in detail. First, we investigated the effect of spine volume-to-surface area. This parameter can be changed in multiple ways – By changing the shape of the spine head and the shape or presence of the spine apparatus (Figure 2 and Figure 3); by changing the size of the spine head alone (Figure 4); and by changing the size of the spine apparatus alone (Figure 5). Next, we investigated the effect of spatial distribution of the fluxes on the PM (Figure 6). Finally, we demonstrate the effect of buffer types and their location (Figures 7 and 8). We describe each of these results in detail below.

**Fig 2.**
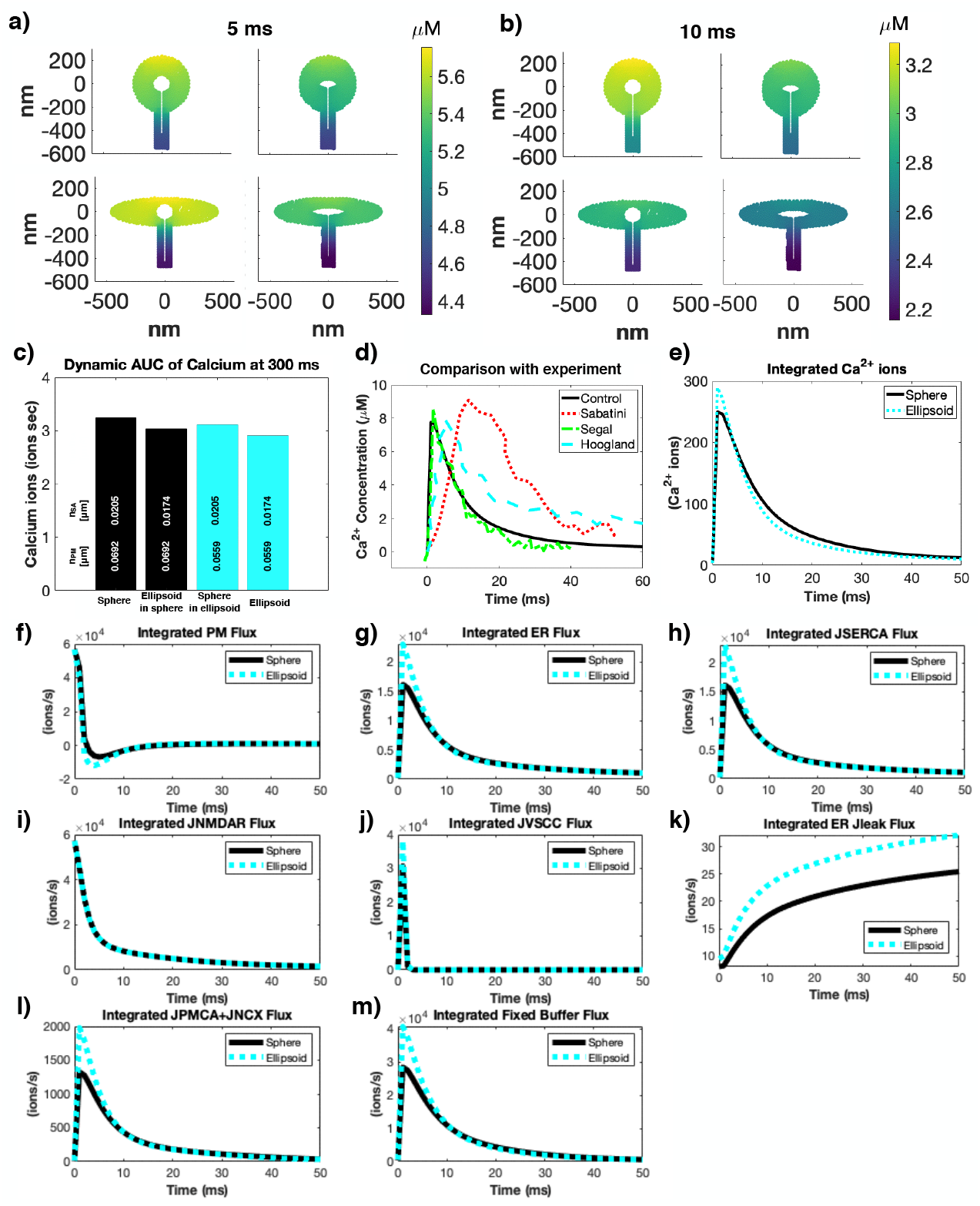
Spine head volume-to-surface area ratio modifies calcium dynamics through membrane flux contributions. a-b) Spatial distribution of calcium in spines at two different time points (5 ms (a) and 10 ms(b)). The instantaneous gradient of calcium ions depends on the shape of the spine head and the shape of the spine apparatus. c) Calcium accumulation at 300 ms was calculated using the area under the curve (AUC) of the spatial and temporal dynamics of calcium throughout the volume; the differences between the shapes are small with the most pronounced difference being a 12% increase in AUC between the sphere and ellipsoid spines. d) We plot the temporal dynamics at the top of the spherical control spine versus reported experimental calcium transients from [27, 63, 64]. The experimental transients are reported in terms of fluorescence which we assume are linearly proportional to concentration [65]. We plot Fig. 1F from [27], Fig. 1 from [63], and Fig. 2D from [64]. We more closely compared the spherical and ellipsoidal spines by integrating total calcium ions over time (e) and considering the integrated fluxes for both shapes (f-m). We see that the ellipsoid has more calcium ions than the sphere (e) because despite having more calcium influx due to VSCC (j), the subsequent higher calcium concentration leads to higher efflux due to pumps (h,l) and buffers (m). Note the timescale in e-m is shortened to 50 ms for clarity.

**Fig 3.**
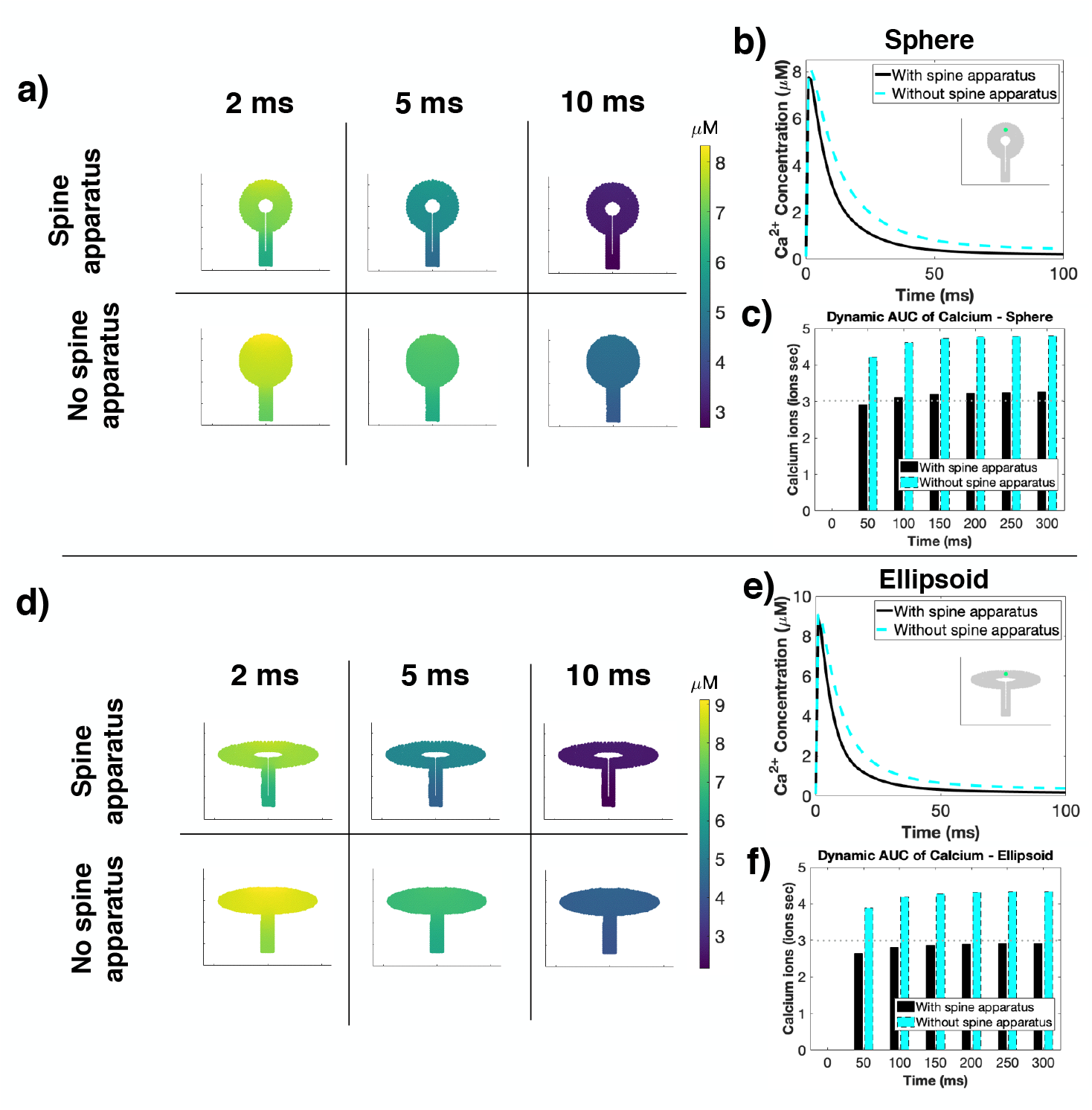
Presence of a spine apparatus, acting as a sink, modulates calcium dynamics. Spines without spine apparatus have higher and more sustained calcium activity despite their increased volume in both spherical (a) and ellipsoidal (d) spines. Temporal dynamics (b,e) and AUC (e,f) plots show that the absence of a spine apparatus (no *J*_*SERCA*_) leads to a prolonged calcium transient and higher total calcium levels for both spherical and ellipsoidal shapes. Inset for b and e show the location in the spine from where the time courses were plotted.

**Fig 4.**
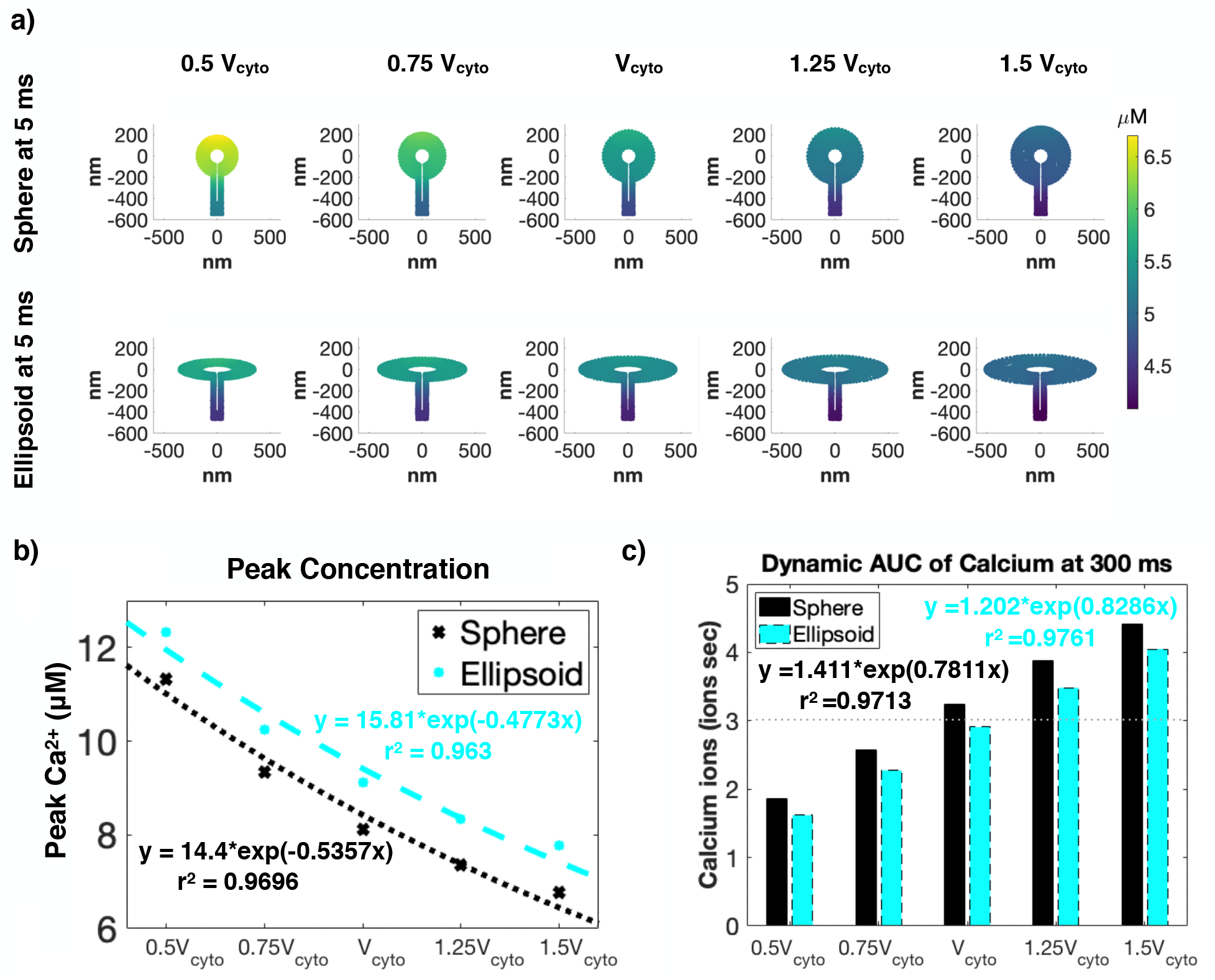
Accumulated calcium scales inversely proportional to the spine head volume. a) Calcium dynamics in spines of different sizes show that as spine volume increases, calcium concentration in the spine decreases. The effect of spine size on the temporal dynamics of calcium is seen in (b) the peak values and (c) AUCs of calcium. Increasing spine volume decreases the peak calcium concentration but increases overall AUC in the spine irrespective of the spine shape.

**Fig 5.**
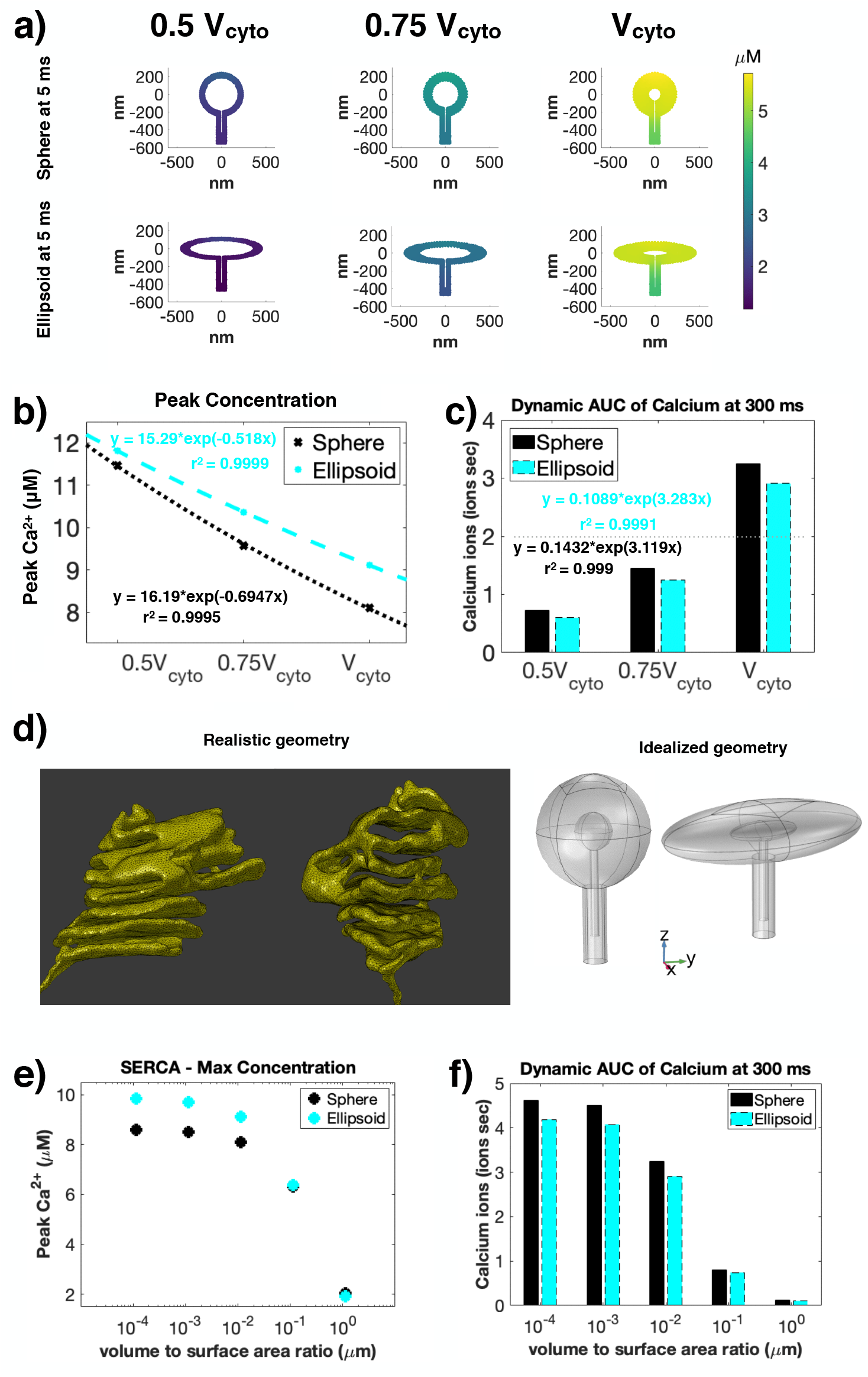
Increasing spine apparatus volume reduces accumulated calcium and spine apparatus volume-to-surface area modulates its ability to act as a sink. a) Calcium dynamics depends on the size of spine apparatus; decreasing cytoplasmic volume by increasing spine apparatus size results in a smaller calcium concentration when compared with a larger spine volume with smaller spine apparatus. The effect of spine apparatus size on the temporal dynamics of calcium is seen in the (b) peak values and (c) AUCs of calcium. Increasing spine apparatus volume (decreasing spine volume) decreases calcium concentration in the spine (a,c), but leads to higher peak concentrations (b). For both geometries, the peak calcium concentration increases for decreasing volume, and can be fit to exponential curves. d) While we are used idealized geometries for both the spine and spine apparatus, in reality the spine apparatus has a complex, helicoidal structure. We investigate this realistic geometry by changing the *n*_*SpApp*_ contribution in the SERCA flux equation. We see that increasing *n*_*SpApp*_ makes SERCA more effective, leading to lower peak concentrations (e) and lower AUC (f). However, we see that as we decrease *n*_*SpApp*_, the change in peak concentration and AUC plateaus, representing highly inefficient SERCA pumps.

**Fig 6.**
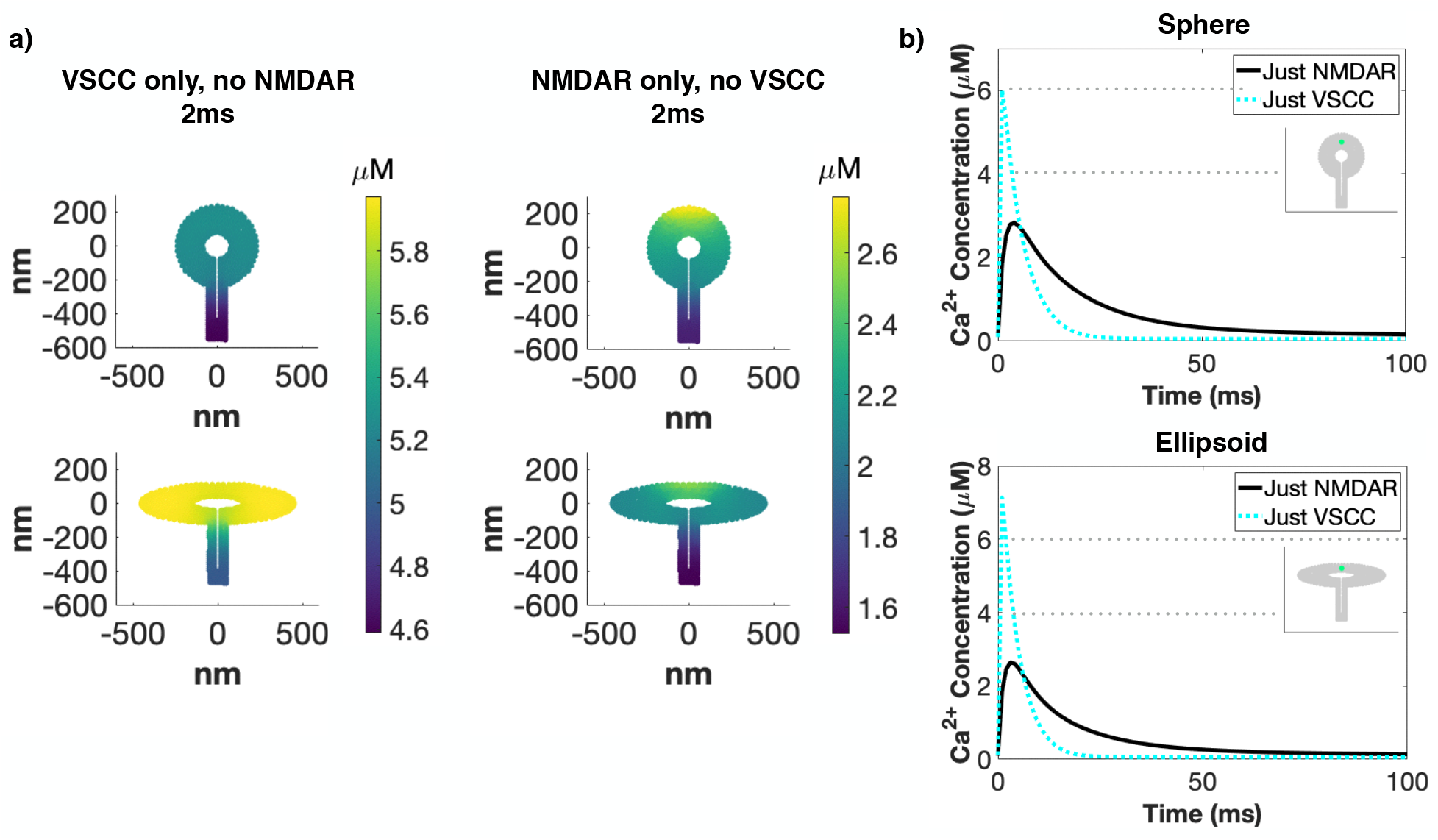
Localization of membrane fluxes alters the spatiotemporal dynamics of calcium. a) Spatial dynamics at 2 ms for spherical and ellipsoidal spines with only one of VSCC or NMDAR as the calcium source. When the main calcium source is VSCC, we see a more uniform concentration with the main gradient between the spine head and spine neck. When NMDAR is the main calcium source, we see a large spatial gradient with a higher concentration at the PSD because the NMDAR is localized to the PSD. b) Temporal dynamics for spherical and ellipsoidal spines with either VSCC or NMDAR as the calcium source. Temporal dynamics clearly show that VSCC act on a faster timescale and have a higher peak calcium when compared to the NMDAR. However, NMDAR influx leads to a more prolonged calcium transient.

**Fig 7.**
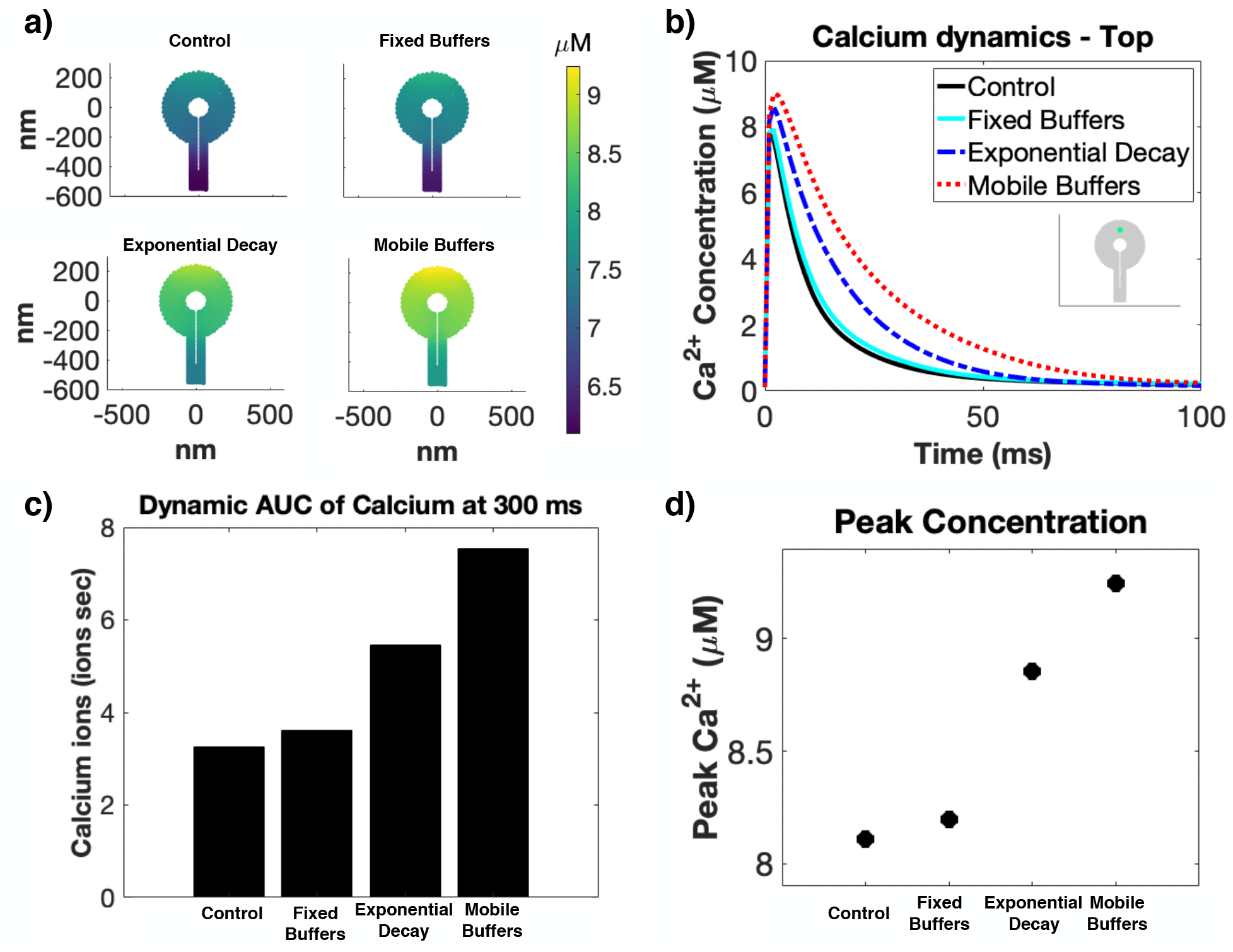
Calcium buffers and CBPs modify all aspects of calcium dynamics. a) Spatial dynamics at 2 ms for spherical spines with different buffer conditions; control (with both fixed and mobile buffers), only fixed buffers, only mobile buffers, and a lumped exponential decay. While all buffer cases show relatively similar peak concentrations (d), all other quantifications show that buffer type greatly impacts the calcium transient decay time (a,b,c). Temporal dynamics (b) show that the control and fixed buffer cases have much faster decay, which translates into lower AUC values (c).

**Fig 8.**
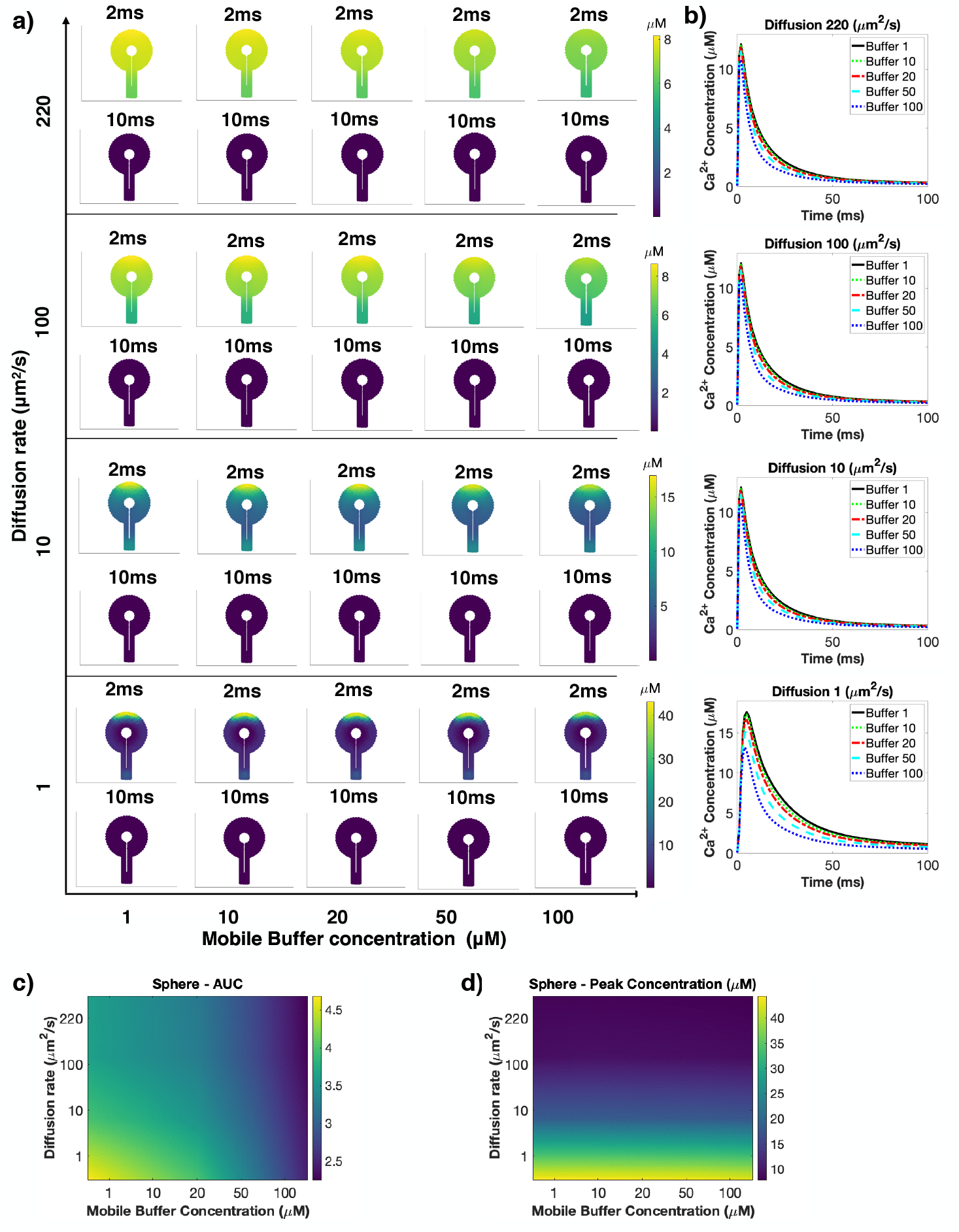
Calcium diffusion rates control spatial gradients of calcium while buffer concentrations control transient decay dynamics. a) Spatial dynamics at 2 ms and 10 ms for a spherical spine. We varied the diffusion coefficient and the mobile buffer concentration. Based on this phase diagram, the diffusion coefficient dictates the range of the spatial gradient of calcium, while buffer binding rate influences the lifetime of the spatial gradient. b) Temporal calcium dynamics at the top of the spherical spine. The temporal dynamics show that the concentration of mobile buffer affects the lifetime of the calcium transient, as expected. c) AUC shows that lower mobile buffer concentration and lower diffusion rates leads to higher levels of total calcium. d) Peak concentration is primarily determined by the diffusion rate of calcium, and is almost independent of mobile buffer concentration.

### Effect of spine volume-to-surface area on calcium dynamics

#### Effects of shape of the spine head and the shape of the spine apparatus on calcium dynamics

We first analyzed how the volume-to-surface area ratio of the spine affects the spatiotemporal dynamics of calcium by simulating calcium dynamics in the different geometries shown in Figure 1c. All of these geometries were constructed such that they have the same volume of the spine cytosol but the surface area of the PM and SpApp vary because of the shape (Table S8). We note that in all the geometries, the temporal dynamics of calcium shows a rapid increase in the first 2-3 ms and a decay over approximately 50 ms (Figure S8). This time course is consistent with experimentally observed and computationally modeled calcium dynamics [42, 57, 58] (Figure 2d and Figure S2, respectively). The spatial profiles of calcium in spheres and ellipsoids (Figure 2a,b) show that spine shape can alter the spatial gradients and decay profiles of calcium. In particular, we observe that while at 5 ms all shapes demonstrate a gradient from the PSD region to the spine neck, because of the localized influx of calcium through the NMDAR in the PSD; at 10 ms, the ellipsoidal spine heads have almost no gradient within the spine head. Instead, they have a more pronounced difference between the spine head and spine neck. We also changed the shape of the spine apparatus to get different combinations of spine head and spine apparatus geometries (Figure 2). Since the apparatus acts as a sink, we find that the reduction in surface area to volume ratio of the spherical apparatus leads to less influx into the spine apparatus, compared to the ellipsoidal spine apparatus (Figure 2c). This result emphasizes the non-intuitive relationships between organelle shape and spine shape.

In addition to the transient response of calcium, the cumulative calcium (total calcium) also carries information with respect to synaptic plasticity [1, 2, 59]. Therefore, we calculated the integrated calcium over the entire spine volume (area under the curve – AUC [60–62]) over 300 ms. Calcium AUC at 300ms (Figure 2c) shows that all spines have slightly different accumulated calcium. Upon closer examination of the membrane fluxes of the sphere and ellipsoid spines (Figure 2e-m), we see that there is a complex relationship between the calcium ion concentration and nonlinear flux terms. In particular, the higher calcium influx in the ellipsoids due to the larger surface area leads to a higher efflux of calcium through the PMCA, NCX, and SERCA pumps and increased binding to fixed buffers on the PM. Therefore, the nonlinear effects of the fluxes associated with the pumps reduces the difference in calcium between spines of difference shapes.

#### Effects of the presence/absence of the spine apparatus on volume-to-surface area

Another way to modulate the volume-to-surface area ratio is to consider spines with and without the spine apparatus. In our model, the spine apparatus serves both as a calcium sink through SERCA pumps and as an excluded volume. We observe that the presence of the spine apparatus in both spheres and ellipsoids (Figure 3a, d top row) results in a steeper gradient from the PSD to the neck when compared to the spines without a spine apparatus (Figure 3a, d bottom row). Additionally, at 10 ms, spines without a spine apparatus have a higher calcium concentration than spines with a spine apparatus. This is because the dynamics of calcium are altered in the presence of the spine apparatus by the SERCA fluxes, Equation (3). As a result, regardless of the shape, spines with a spine apparatus have a faster decay of cytosolic calcium (Figure 3b, e). We found that in both geometries, the AUC of spines with a spine apparatus was lower at different time points when compared to spines without a spine apparatus (Figure 3c, f). At 300 ms, the spine without a spine apparatus has 47.75% more total calcium for the spherical geometry and 49.09% more for the ellipsoidal geometry. Thus the flux due to SERCA plays a significant role in altering the decay dynamics of calcium in spines with spine apparatus, and as a result alters the total calcium.

### Effect of spine volume on calcium dynamics

In addition to spine shape, spine size (volume) is also known to change during development and plasticity related events [1, 66, 67]. How does changing the volume of the spine, while maintaining the same shape affect calcium dynamics? To answer this question, we conducted simulations in spherical and ellipsoidal spines of different volumes. Recall that the control spine has a cytosolic volume *V*_*cyto*_ 0.06*µm*^3^ (see Table S8). For each geometry (sphere and ellipsoid), we maintained the same size and shape of the spine apparatus as before and only changed the spine cytoplasm volume in the range of 0.5*V*_*cyto*_ to 1.5*V*_*cyto*_, (Table S9 and Table S10). We found that the relationship between spine volume and calcium concentration is inversely proportional. For spine volumes smaller than the control, we observed an increase in calcium concentration for both geometries, whereas for larger volumes, calcium concentration decreases (Figure 4a,b). As expected, we found that for both geometries, an increase in spine volume resulted in an increase in cumulative calcium, (Figure 4c). Furthermore, we found that the change in cumulative calcium has a direct but nonlinear relationship with the change in spine volume. For the range of volumes investigated, the peak calcium concentration and AUC show an exponential relationship with respect to volume. We see at all sizes the ellipsoid has higher peak concentrations but lower AUC compared to the sphere.

### Effect of spine apparatus size and geometry

The complex architecture of the spine apparatus was recently elucidated in a focused ion beam scanning electron microscopy study by Wu et al. [24]. Since we cannot yet manipulate the shape of the spine apparatus *in vivo*, we varied the geometric features of the spine apparatus *in silico* to see how they affect calcium dynamics. Previously, we showed that spine apparatus shape and spine volume separately can alter the AUC and peak calcium. Here, for a given spine shape, we varied the volume of the spine apparatus to modulate the cytosolic volume to spine apparatus volume ratio. In this case, by varying the spine apparatus volume, we altered the spine volume to be 50% and 75% of V_cyto_, the control spine volume.

Here, we change the spine volume by the changing spine apparatus size (Figure 5). We found that a larger spine apparatus leads to an decrease in calcium concentrations and AUC (Figure 5a,c), but an increase in peak concentration (Figure 5b). We note that as spine apparatus volume increases, AUC drops in a nonlinear manner in both geometries. The spherical spine has a 55.4% and 77.8% reduction in AUC from control for the 75% and 50% spines, while the ellipsoidal spine has a 57.2% and 79.4% reduction from control. For all sizes, the ellipsoid shows higher peak concentrations but lower AUC compared to the sphere. This effect is in part due to the shortened distance between the PSD and spine apparatus in the ellipsoid. The distance between the PSD and spine apparatus is an important lengthscale in the spine since it controls the distance between the sources and sinks of calcium. Therefore, changing this lengthscale, as happens when changing the spine apparatus volume, influences calcium dynamics in nonlinear ways. From these observations, we conclude that spine volume coupled with spine apparatus volume and surface area is an important regulator of calcium dynamics in dendritic spines.

An intriguing feature of the spine apparatus, in particular (Figure 5d) [24], and the ER, in general [68], is the large surface-to-volume ratio occupied by this organelle. Therefore, we next considered the effect of the volume-to-surface area ratio ‘n’ (given in units of length) of the spine apparatus (Figure 5d-f). We modeled the boundary flux on the SpApp membrane such that this flux is proportional to n_SpApp_ (volume-to-surface area ratio for the spine apparatus). As a result, when we increase the ‘volume’ of the spine apparatus by increasing n_SpApp_, calcium flux into the SpApp will increase. We noticed that at lower n_SpApp_ values, the peak calcium concentration and to a less obvious extent the AUC (Figure 5e-f) plateau, but decrease substantially at larger n_SpApp_ values.

From these observations, we conclude that the spine apparatus acts as a physical and spatial buffer for calcium dynamics by regulating the timescale through surface to volume regulation in the interior of the spine. The SpApp acts as a calcium sink in the timescale of interest [31, 50, 51] and in the absence of the SpApp, the only way to remove calcium from this system is through CBPs and pumps. Furthermore, since the SpApp has been known to grow and retract from the dendritic spine in response to stimuli [69], regulation of SpApp surface area can also allow for rescaling calcium dynamics in the spine [70].

### Spatial distribution of membrane fluxes governs calcium dynamics supralinearly in dendritic spines

The density of VSCC and number of NMDAR that open in response to stimuli in dendritic spines varies [71]. One of the primary determinants of calcium dynamics in the spine is the various membrane fluxes because these fluxes serve as sources and sinks at the PM and sinks at the spine apparatus membrane. We investigated the effect of the spatial distribution of membrane fluxes on calcium dynamics in the dendritic spine, by considering either only NMDAR or VSCC as the calcium source (Figure 6). We observed that if only NMDAR activity was present, then calcium concentration was high in the PSD region due to the localization of NMDAR to the PSD but the overall calcium concentration was small regardless of the spine head shape (Figure 6a). However, if only VSCCs were active, then the spatial gradient of calcium is mainly between the spine head and spine neck (Figure 6a). The temporal dynamics are also affected by the receptor and channel distributions (Figure 6b). With only VSCC present, there is a larger calcium peak but with faster decay. With only NMDAR, we observe a lower calcium peak concentration but a prolonged transient. Therefore, membrane flux distribution can impact the spatial and temporal dynamics of calcium in a nonlinear manner. This agrees with experimental results stating that the various calcium sources behave supralinearly [16] and a balance between various calcium fluxes is required for tightly regulating calcium concentrations in these small volumes.

### Calcium buffers and diffusion couple to alter calcium spatiotemporal dynamics

Due to the vast number of buffers that are known to affect calcium dynamics, we investigated the effect of four different buffer conditions – i) both fixed membrane-bound buffers and mobile cytosolic buffers (control), ii) fixed buffers localized to the membrane, iii) mobile buffers in the cytoplasm, and iv) a uniform exponential decay applied across the whole cytoplasm (Figure 7 and SOM Figure S9). We observe similar spatial dynamics for all buffer types (Figure 7a), but temporal dynamics differ greatly (Figure 7b) and as a result alter accumulated calcium (Figure 7c). We see that the decay behavior of calcium, and therefore total calcium, is highly dependent on buffer type. Peak calcium follows the same trend as AUC, with mobile buffers having the highest calcium concentrations and total calcium, then exponential decay, fixed buffers, and finally the control. We also consider how reaction dynamics versus diffusion rate govern calcium dynamics because buffer dynamics and diffusion rates of calcium are coupled [10]. We varied the diffusion rate of calcium and the concentration of mobile buffers to quantify how calcium dynamics are reaction- or diffusion-controlled (Figure 8). We see that diffusion controls the spatial gradient seen within the spine (Figure 8a), while mobile buffer concentration controls the lifetime of the calcium transient (Figure 8b). Combining these effects, we see that the buffer concentration variation has the greatest effect at lower diffusion rates (bottom row of Figure 8a,b, and Figure 8c). The peak concentration of calcium is almost entirely dependent on the diffusion rate (Figure 8d). Therefore, based on the high diffusion rate of Ca^2+^ reported in the literature, we expect the system to be in a diffusion-dominated regime.

## Discussion

Calcium is a fundamental player in both neuronal (cellular) and neural (systems) functionality [72–74]. Compartmentalized by dendritic spines, calcium has a vital role in triggering signaling pathways for LTP, LTD, synaptic plasticity, and other processes associated with learning and memory formation [1]. However while dendritic spines are known to form functional subcompartments, it is less understood how this specialized compartmentalization impacts calcium dynamics [10]. In this study, we explored the intricate relationship between calcium dynamics and the shape and size of dendritic spine structures. We found that while the relationship between spine geometry and calcium dynamics is quite complicated [29, 30, 75], some general conclusions can be drawn from our study.

First, the volume-to-surface ratio, rather than the shape and size itself seem to have a dramatic effect on spine calcium (Figures 2 to 5). Of course, the volume-to-surface ratio itself can be dramatically altered by size, shape, and internal organization as a many-to-one function. Then, we can think of the ultrastructural organization of the dendritic spine [34] as perhaps ‘optimized’ to not just increase contacts with neighboring axons and neural circuit connection [10, 16, 74] but also to tune this volume-to-surface ratio dynamically [76]. This volume-to-surface ratio coupling further highlights the complex relationship between spatial sources and sinks of calcium, which becomes apparent when the distance between the spine plasma membrane and internal organelle becomes quite small. We note that in our model we assume constant pump density, which highlights the volume-to-surface area ratios between various shapes [77, 78]. Experimental results have already shown different behavior in large versus small dendritic spines [56], and additional studies on dendritic spine geometry have shown that stable, mature spines tend to be larger spines that tend towards mushroom shapes as they grow around adjoining axons, and are more likely to have a spine apparatus [34]. In comparison, younger, less stable spines tend to be smaller and more spherical [30, 34]. Therefore, we predict that spine size and spine apparatus presence are coupled to control calcium dynamics. This result should be investigated further, in particular to make predictions on why stable spines tend to be larger and mushroom shaped. The inverse relationship between the volume-to-surface ratio that we found, and a possible exponential relationship suggest a possible limiting mechanism for maintaining homeostasis of synaptic potentiation [79–82]. By altering the spine size dynamically [76] and the presence and absence of the spine apparatus dynamically [69], spines could maintain their optimal range of synaptic function [79–82].

Second, localization of membrane fluxes alters calcium transients (Figures 5 and 6). These fluxes, which serve as boundary conditions, can be altered by changing the density and distribution of calcium sources and sinks. This idea is consistent with how calcium signal localization is a result of tuning the distance between sources and sinks [8]. We show here that in addition to distance, the strength of the fluxes is important. Thus, in various disease states that impact the distribution or strength of membrane components, we predict atypical spatial calcium gradients are possible that could impact downstream signaling pathways. For example, NMDAR dysfunction whether leading to increased or decreased functionality can potentially lead to central nervous system diseases such as stroke, Huntington’s disease, or schizophrenia [83].

Finally, the role buffers play in modulating calcium transients is not just by changing the decay time as previously thought, but also by tuning the membrane fluxes, especially in the case of fixed buffers (Figures 7 and 8) [22, 49]. Again, the timescale that we see is a combination of rate alterations at the membrane and rate alterations in the volume resulting in broader control of calcium dynamics. The crowded environment within the spine head also has consequences for calcium diffusion and while it is possible for calcium to diffuse through a crowded space, particularly in the PSD, the exact mechanisms of such transport remain unclear [38, 84, 85]. Thus, our study highlights the need for connecting biophysical features of the spine and molecule localization to the dynamics of calcium (Figure 1).

We also note that our model has certain limitations. In particular, within the crowded environment of the dendritic spine cytoplasm is an abundance of actin, which has previously been shown to have the potential to create cytosolic flow through contraction following spine activation, leading to faster calcium diffusion [20]. We do not address this spine contraction in this model, but actin contributions are a focus of ongoing research in our group. While we touched upon the role of diffusion and CBPs, we also acknowledge that much work remains to be done on the true impact of the dense actin network and crowded environment within dendritic spines in regulating these processes [86]. In addition, we modeled isolated spines, but the width of the spine neck has been showed as an important determinant of calcium dynamics when comparing larger and smaller spines [43, 77, 87] and could play into communication to the dendrite. It is also possible that stochastic modeling will give better quantitative insights without altering the underlying physics [6, 36, 88]. The development of a combined stochastic-deterministic model can help combine these two regimes to address the fundamental physics that occurs in these small systems with complex membrane dynamics and few molecule situations.

Despite these shortcomings, we have identified several key features of the relationship between dendritic spine geometry and calcium dynamics. Current models of synaptic weight updates use calcium as the determinant for the synaptic weight vector and the learning rate [13, 89–91]. Here, we show that calcium in a spine, even in short timescales, is a function of geometry, ultrastructure, and buffers. Based on these insights, we speculate on what the biophysical features of the spine mean for neural systems-level functionality. It has long been considered that a neural circuit level model of synaptic weight updates can be informed by the calcium transients in the synapse. As a result, weighting functions have been proposed that consider calcium dynamics [13, 89, 90] and these functions have been used to connect biophysical features of NMDAR to synaptic weight updates in models of STDP [92–96]. We now propose that the calcium transient is *explicitly* a function of the spine volume-to-surface area, ultrastructure, and buffers, and that the calcium functions that inform the synaptic weight vectors and synaptic learning rate (Figure 9) must be updated to consider such geometric information. We anticipate that such new models can give us better insight into the neural circuitry of the brain and also better inform bioinspired engineering of neuromorphic circuits [97]. We also acknowledge that much work remains to be done in connecting the spatial signaling aspects in postsynaptic spines with neural circuit behavior but hope that this work will inspire more multiscale modeling efforts in this field. The spatial aspects of calcium dynamics are also fundamental towards understanding the downstream dynamics of critical molecules such as CaMKII, the small RhoGTPases (Cdc42, Rho, and Rac), and subsequently actin dynamics in dendritic spine remodeling [1, 4, 76, 98–102]. Going beyond single spine dynamics, the propagation of the downstream mechanochemical activity to neighboring spines is a key step towards integrating single spine behavior to multiple spines, across the dendrite, and ultimately the whole cell [4, 7, 103–105]. Thus, we posit that accounting for the spatial and physical aspects of calcium dynamics is the first step towards deciphering the complex shape function relationships of dendritic spines.

**Fig 9.**
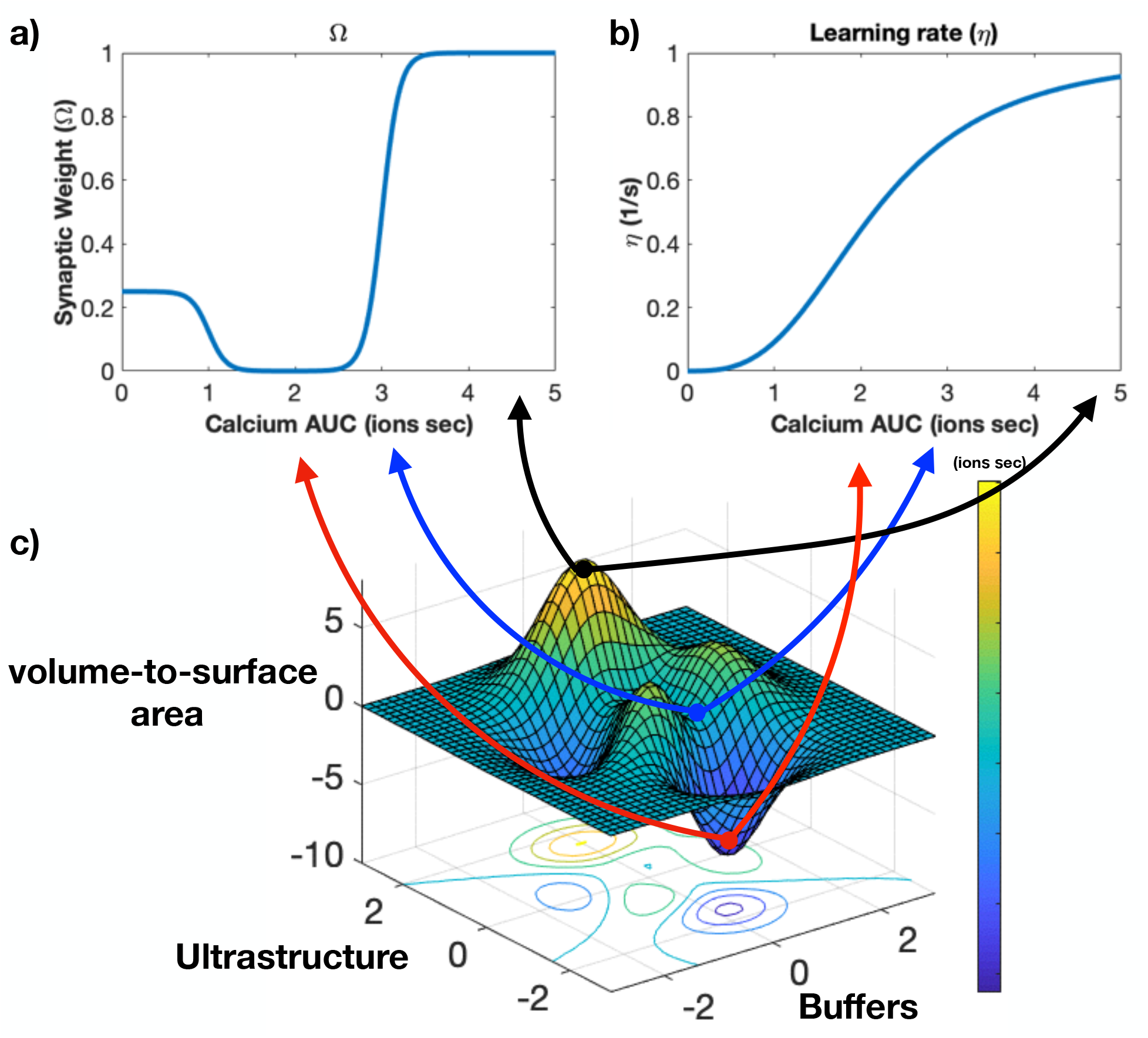
Biophysical factors can impact synaptic weights through calcium dynamics. a) Synaptic weight can be calculated from calcium dynamics [13]. We plot changes in synaptic weight due to calcium quantified through accumulated calcium. b) Calcium dynamics also dictate the learning rate of the spine. c) Qualitative representation of the effects of calcium AUC on synaptic weight and learning rate. Using our model, we can map how various factors governing calcium dynamics influence both synaptic weights and learning rates of dendritic spines. This surface plot visualizes how three different factors, volume-to-surface area ratio, ultrastructure, and calcium buffers couple to influence calcium AUC (colorbar) that feeds back into synaptic weight changes and learning rates.

## Abbreviations

*Ca*^2+^: Calcium
PM: Plasma membrane
ECS: Extracellular space
ER: endoplasmic reticulum
SpApp: spine apparatus
cyto: cytoplasm
AUC: area under the curve
CBP: Calcium binding proteins.

## Acknowledgments

We thank various members of the Rangamani Lab for valuable discussion, Mariam Ordyan for manuscript review, and Professor Brenda Bloodgood and Evan Campbell for their expertise. We thank Professor Pietro De Camilli and his laboratory for supplying images used to construct the mesh seen in Figure 5d. We would also like to thank the UCSD Interfaces Graduate Training Program and the San Diego Fellowship for funding and support. MB was also supported by AFOSR MURI grant number FA9550-18-1-0051 to PR and TJS and a grant from the National Institutes of Health, USA (NIH Grant T32EB009380).

## S1 Model development and analysis

The spatial dynamics of calcium can be modeled as a reaction-diffusion equation in the volumetric space between the plasma membrane (PM) and the spine apparatus (SpApp) membrane in the spine, where the primary reaction is the binding of calcium by calcium binding proteins (CBPs) in the cytosol. The governing equation is given by

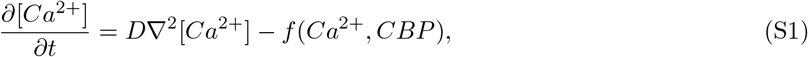

where *f* (*Ca*^2+^*, CBP*) represents the rate at which the cytosolic calcium reacts with the CBPs. It is well known that solutions to PDEs such as that in Equation (S1) are specific to the prescribed boundary conditions [106]. The corresponding boundary conditions on the PM are an inward flux of calcium due to the NMDAR and VSCC activation and an outward leak due to the PM pumps. We also model calcium binding to fixed buffers on the plasma membrane as an outward flux onto the membrane. At the PM, the boundary condition can be written as

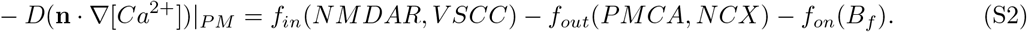

Similarly at the spine apparatus membrane, the boundary condition accounts for the action of SERCA pumps pulling calcium into the spine apparatus and a leak from the spine apparatus into the cytosol, given by

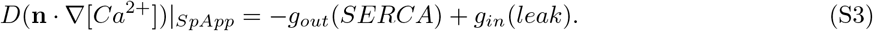

The analytical solution to similar equations as Equation (S1) for linear boundary conditions was described in detail for ellipses and circles in [106]. Additional analytical and numerical analyses of a similar system with time dependent boundary conditions can be found in [55]. Here, we extend this analysis to the non-linear, time-dependent boundary conditions associated with the calcium receptors, channels, and pumps.

### S1.1 A comment on length scale

When converting between an ordinary differential equation model and a partial differential equation with boundary conditions, volume equations can become boundary fluxes. In this case, we need to use a length scale (n [*µ*m]) to convert from volume units to the appropriate flux units. In our model, we use 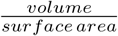 of the respective compartment, ex. cytoplasm or spine apparatus. However, to have the most realistic length scale possible, we utilize *n*_*r*_ which is a length scale taken from the corresponding compartment in a realistic geometry [107]. Based on how we define various fluxes, we use this conversion for the various PM and SpApp pumps (*J_PMCA_*, *J*_*NCX*_, *J*_*SERCA*_), and the SpApp leak term (*J*_*LEAK*_). *J_VSCC_* is always defined as a constant density of channels and *J*_*NMDAR*_ is defined to have a constant total flux (integrated flux). We should note that by using a constant length scale ‘n,’ we are effectively defining a constant density of flux. We can compare this with the case in which we use a specific length scale such that ‘n’ is the ratio of the volume-to-surface area of the specific geometry in the model (Figure S1). In the case of a specialized n conversion length scale, we conserve volume across shapes but the surface area changes. However, the nonlinearities of the pump flux terms coupled with the calcium influx due to VSCC being surface area-dependent complicate the effects of choice of ‘n.’

**Fig S1.**
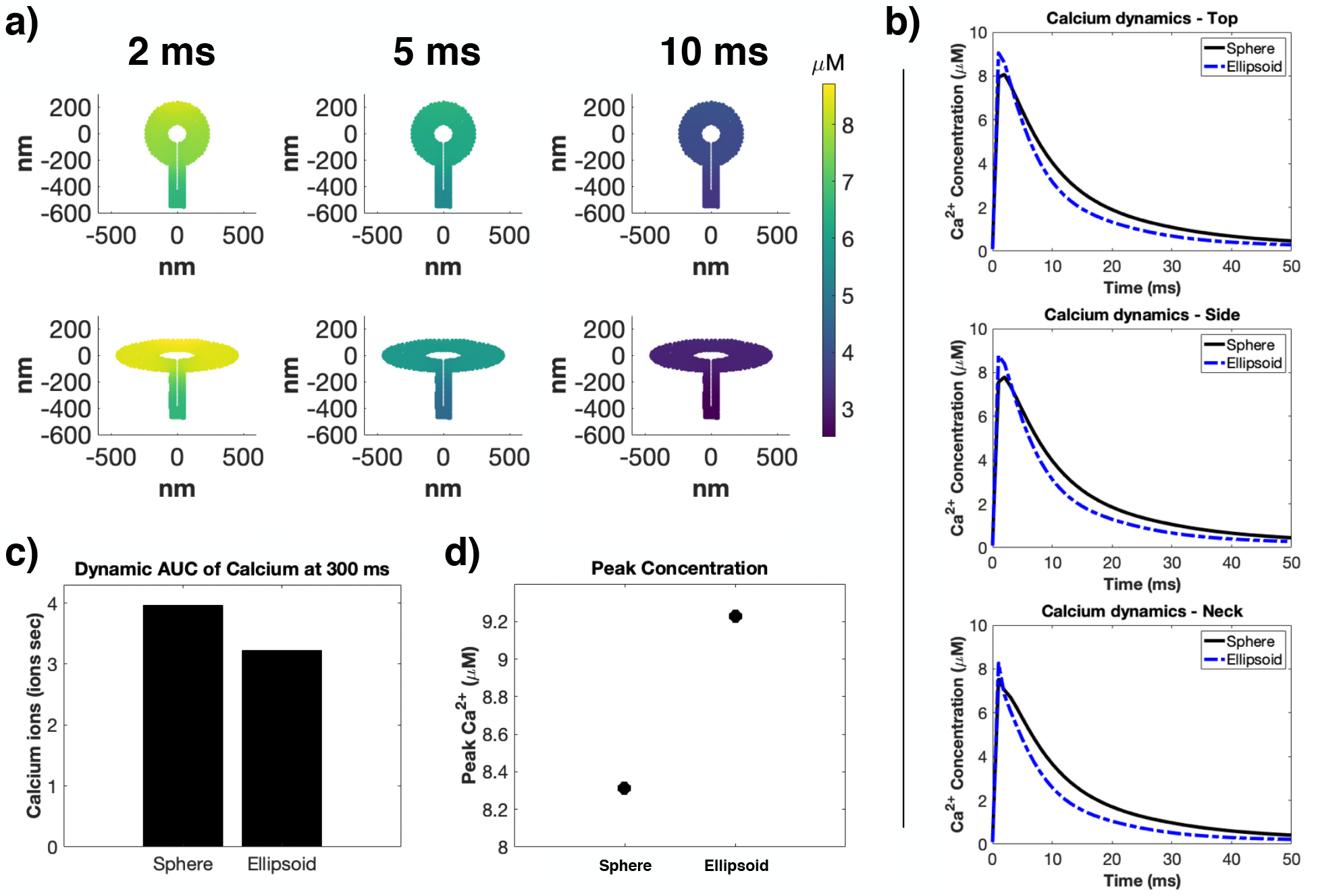
Variable pump density exaggerates volume-to-surface area differences. For our model, we assume constant pump density through our choice of the ‘n’ length scale. However, we can use specific ‘n’ for the geometry of a specific simulation’s geometry. Here we use geometry-specific ‘n’ for all pump and leak terms. a) Spatial profiles of calcium concentration at three time points (2ms, 5ms, and 10ms) for both spherical and ellipsoidal control spines. b) Temporal calcium dynamics for the spherical and ellipsoidal controls at three spatial locations (top, side, and neck). c) Calcium AUC at 300 ms for both control spines. d) Peak concentrations for both control spines.

### S1.2 Calcium pumps and channels

There are 2 volumes (cytosol and spine apparatus) and two boundaries (plasma membrane and spine apparatus membrane) of interest. We assumed that the extracellular space (ECS) acts as a constant reservoir of calcium due to its high basal calcium concentration. Calcium dynamics are governed by partial differential equations (PDEs) and their accompanying boundary PDEs. The main equations in the model are summarized in Table S1.

#### S1.2.1 Volume

The cytoplasm has simple diffusion and decay from CBPs that are present in the cytosol. We investigated four different forms of CBP - 1. exponential decay; 2. immobile/fixed buffers; 3. mobile buffers; and 4. immobile/fixed and mobile buffers. We have a general form of

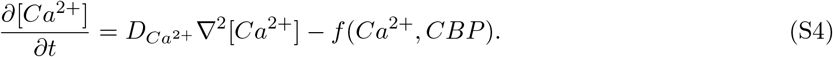

**Table S1.**
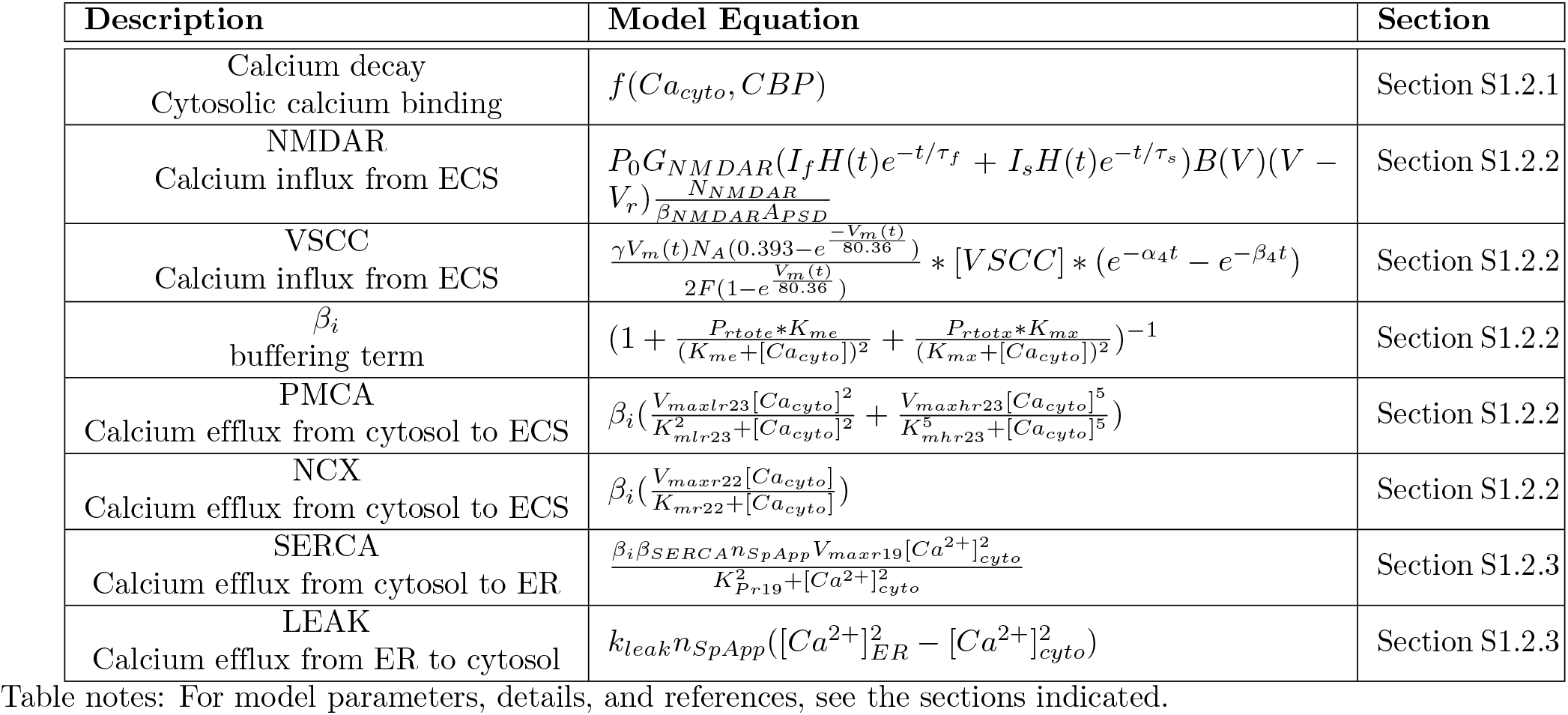
Main equations in the model.

For simple exponential decay, the equation becomes

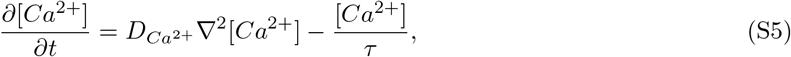

where *τ* is a decay time constant. *τ* is picked to match experimentally observed and computationally modelled calcium decay dynamics [42]. We plot the temporal dynamics at the top of the spherical control spine versus temporal dynamics reported in both computational (Figure S2) and experimental (Figure 2d) calcium studies.

The identities of immobile/fixed buffers have remained fairly elusive, but there has been some speculation that fixed buffers are near the membrane, potentially as receptors and channels that bind calcium. Therefore, we localize the fixed buffers to the membrane. This transforms a volume reaction to a boundary flux. Calcium binds with our fixed buffers on the membrane, represented as a flux boundary condition onto the membrane. Therefore, cytosolic dynamics are simply

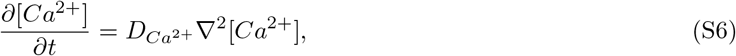

with an additional boundary condition representing fixed buffers binding calcium from the cytoplasm given as

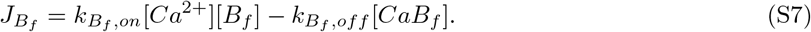

The fixed buffer and calcium bound fixed buffer have reactions defined as

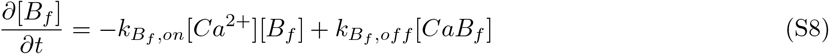

and

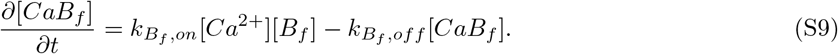

Mobile buffers are modeled as a diffusive species in the cytoplasm, so the cytosolic species dynamics are given by

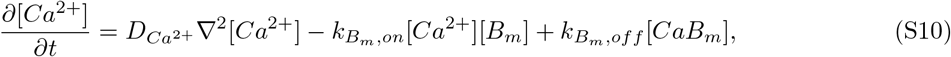

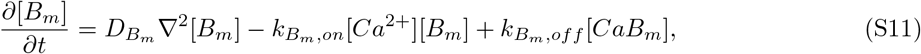

and

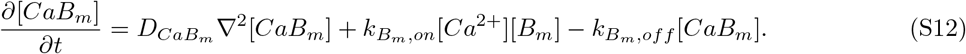

**Table S2.** Parameters used in the model for the volumes.

| Variable | Value | Units | Reference |
| --- | --- | --- | --- |
| $[Ca^{2+}]_{cyto}^{IC}$ | 0.1 | $\mu M$ | [6, 109] |
| $[Ca^{2+}]_{ER}^{IC}$ | 60 | $\mu M$ | [6] |
| $[Ca^{2+}]_{ECS}^{IC}$ | 2 | mM | [110] |
| $\tau_{cyto}$ | 20 | ms | * |
| $D_{Ca^{2+}}$ | 220 | $\frac{\mu m^2}{s}$ | [41, 111] |
| $k_{B_f, on}$ | 1 | $\frac{1}{\mu M \cdot s}$ | [6] |
| $k_{B_f, off}$ | 2 | $\frac{1}{s}$ | [6] |
| $k_{B_m, on}$ | 1 | $\frac{1}{\mu M \cdot s}$ | [112] |
| $k_{B_m, off}$ | 1 | $\frac{1}{s}$ | [112] |
| $D_{B_m}$ | 20 | $\frac{\mu m^2}{s}$ | [41] |
| $D_{CaB_m}$ | 20 | $\frac{\mu m^2}{s}$ | [41] |
| $B_{fIC}$ | 20 | $\mu M$ | [112] |
| $B_{mIC}$ | $78.7 \cdot n_{PM_r}$ | $(\mu M) \cdot (m)$ | [6] |

Table notes: \*This value was estimated in this work to match experimental traces, but was independently verified by [42].

For the cytosol, we set the initial concentration of 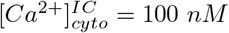. Calcium is a commonly used second messenger and is rarely found unbound in neurons [1, 6, 10, 11, 108]. For the spine apparatus, we assume an initial concentration of 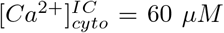. Various studies have found a calcium range between 60 to 200 *µM* in the spine apparatus [6, 39]. For the extracellular space, we set an initial concentration of 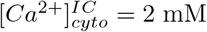 [6, 17].

#### S1.2.2 Plasma Membrane

The plasma membrane (PM) has NMDA receptors (NMDAR), voltage sensitive calcium channels (VSCC), and various calcium influencing pumps (PMCA and NCX). We also have fixed buffers localized to the PM. The effective flux of Ca^2+^ due to these sources can be written as

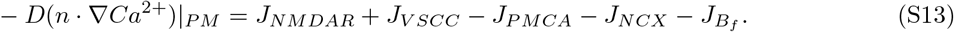

##### Flux due to NMDAR

NMDAR are present in the post-synaptic density (PSD) and open in response to depolarization of the membrane [12, 13]. Therefore, we designate NMDAR as a localized flux at the top of the spine head that acts over the area, *A_PSD_*.

Single channel conductance of NMDAR is given by

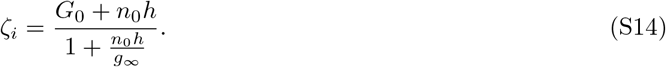

**Table S3.** Parameters for NMDAR.

| Variable | Value | Units | Reference |
| --- | --- | --- | --- |
| $N_{NMDAR}$ | 1 | receptor | [113] |
| $A_{psd}$ | variable | $\mu m^2$ | geometry-dependent |
| $G_0$ | 65.6 | pS | [12] |
| $P_0$ | 0.5 | | [12] |
| $h$ | 11.3 | pS/mM | [12] |
| $g_\infty$ | 15.2 | pS | [12] |
| $r$ | 0.5 | | [110] |
| $K_m$ | 0.092 | $mV^{-1}$ | [14] |
| [Mg] | 1 | mM | [12] |
| $Mg_{scale}$ | 3.57 | mM | [14] |
| $V_r$ | 90 | mV | [114] |
| $I_f$ | 0.5 | | [13] |
| $I_s$ | 0.5 | | [13] |
| $\tau_f$ | 50 | ms | [13] |
| $\tau_s$ | 200 | ms | [13] |
| $\beta$ | 85 | | estimated * |

Table notes: \* This parameter is similar to the parameter $\alpha$ in [12, 13].

*G*_0_ and *g_∞_* are the limiting conductances for low and high *Ca*^2+^ levels, h sets how fast the limiting cases are reached, and *n*_0_ is based on extracellular Ca^2+^ activity. More specifically,

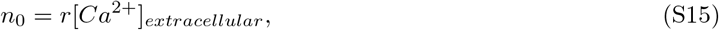

where r is 0.5 given by [110].

Single conductance is given by *ζ_i_*, defined above with units of [S]. We note that one Ca^2+^ ion has two charged particles so the charge carried by this ion is

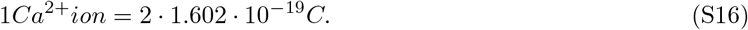

We divide the single conductance by this ion to coulomb ratio and by Avogadro’s number to get units of moles,

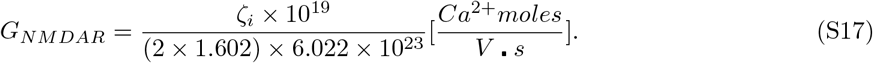

The flux of calcium depends on this conductance and the dynamics of receptor opening. The flux is given by

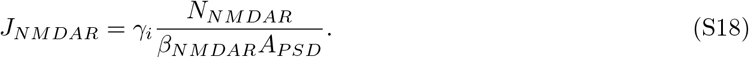

where *β* is a constant,

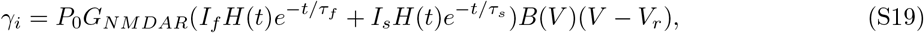

and

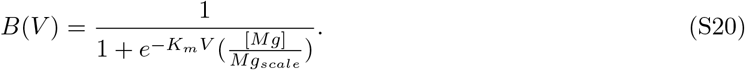

H(t) is the heaviside function. *N*_*NMDAR*_ is the number of open receptors. All parameters for the NMDAR flux are given in Table S3.

**Table S4.** Parameters for membrane voltage.

| Variable | Value | Units | Reference |
| --- | --- | --- | --- |
| Vrest | -65 | mV | [6] |
| maxBPAP | 38 | mV | estimated # |
| tdelaybp | 2 | ms | * |
| $I_{bsf}$ | 0.75 | | [13] |
| $I_{bss}$ | 0.25 | | [13] |
| $t_{bsf}$ | 3 | ms | [13] |
| $t_{bss}$ | 25 | ms | [13] |
| $s_{term}$ | 25 | mV | estimated # |
| tdelay | 0 | ms |  |
| tep1 | 50 | ms | [13] |
| tep2 | 5 | ms | [13] |

Table notes: \* This parameter was chosen to give a maximum possible depolarization based on STDP [41,115,116]. # These parameters based on [12,13] were modified to provide a physiologically relevant membrane voltage depolarization for dendritic spines.

Membrane voltage is time-dependent and depends on the back propagating action potential (BPAP) and excitatory postsynaptic potential (EPSP) [12–14, 42]. Membrane voltage is given by

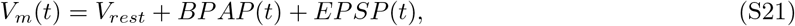

where

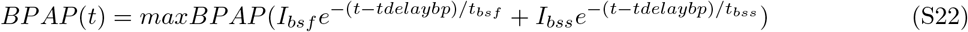

and

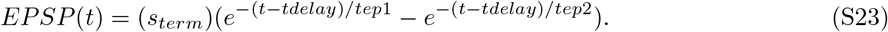

Parameters for membrane voltage can be found in Table S4. We based this input on spike time-dependent plasticity (STDP) such that we get the maximum depolarization [41,115,116]. We estimated several parameters to achieve a physiologically relevant membrane voltage depolarization for a dendritic spine.

##### Flux due to VSCC

We include voltage sensitive calcium channels (VSCC) across the whole plasma membrane. [6] gives the rate of calcium through VSCC as

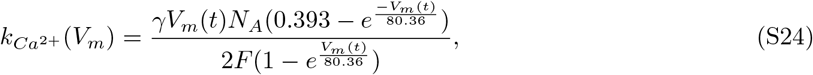

which has units of [1/s]. It also depends on the number of activated VSCC which is a function of Vm. However, we will model this as VSCC multiplied by a time dependent biexponential (based on [1]). Thus, we get

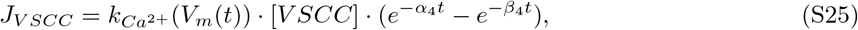

where parameter values can be found in Table S5.

##### Flux due to PMCA and NCX

The plasma membrane has several pumps that work to pump calcium from the cytosol into the extracellular space. We include plasma membrane Ca2+ ATPase (PMCA) and sodium-calcium exchanger (NCX) pumps on all of the plasma membrane of our model. We based the majority of the PMCA and NCX equations on [53, 54, 117], in particular the magnitude of the pump fluxes were compared to those in [117].

**Table S5.** Parameters for VSCC.

| Variable | Value | Units | Reference |
| --- | --- | --- | --- |
| $N_A$ | $6.022 \times 10^{23}$ | ions/mole | Avogadro's number |
| $A_{membrane}$ | variable | $\mu m^2$ | geometry dependent |
| $\gamma$ | 3.72 | pS | [6] |
| F | 96485.332 | C/mole | Faraday's number |
| $\alpha_4$ | 34.7 | $ms^{-1}$ | [6] |
| $\beta_4$ | 3.68 | $ms^{-1}$ | [6] |
| [VSCC] | $2/(6.022 \times 10^{23})$ | $moles/\mu m^2$ | [6] |

**Table S6.** Parameters for PMCA and NCX pumps.

| Variable | Value | Units | Reference |
| --- | --- | --- | --- |
| $P_{rtote}$ | $1.91 \cdot 10^2$ | $\mu M$ | [54] |
| $K_{me}$ | 2.43 | $\mu M$ | [54] |
| $P_{rtotx}$ | 8.77 | $\mu M$ | [54] |
| $K_{mx}$ | 0.139 | $\mu M$ | [54] |
| $V_{maxlr23}$ | 0.113 | $\mu M/s$ | [54] |
| $K_{mlr23}$ | 0.442 | $\mu M$ | [54] |
| $V_{maxhr23}$ | 0.59 | $\mu M/s$ | [54] |
| $K_{mhr23}$ | 0.442 | $\mu M$ | [54] |
| $V_{maxr22}$ | 0.1 | $\mu M/s$ | [54] |
| $K_{mr22}$ | 1 | $\mu M$ | [53] |
| $\beta_{PMCA}$ | 100 | | estimated * |
| $\beta_{NCX}$ | 1000 | | estimated * |
| $n_{PM_r}$ | 0.1011 | $\mu m$ | [107] |
| $vol_{cyto_r}$ | 0.67 | $\mu m^3$ | [107] |
| $area_{PM_r}$ | 6.63 | $\mu m^2$ | [107] |

Table notes: \* These values are used to obtain fluxes comparable to [117].

The calcium buffering term that affects the pumps is given by

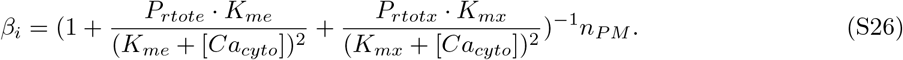

PMCA dynamics are given by

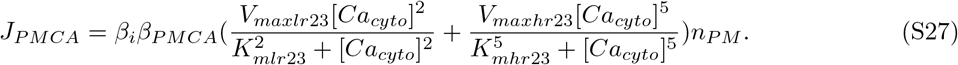

NCX dynamics are given by

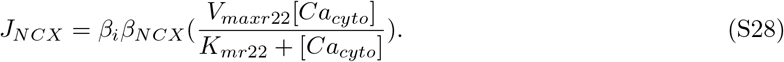

To ensure a membrane flux, we multiply the above equations by the correction factor *n_PMr_* given by

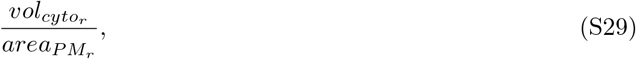

which is the ratio of the spine volume to the surface area of the plasma membrane in a realistic dendritic spine geometry [107]. All parameters for the PMCA and NCX pumps can be found in Table S6.

**Table S7.**
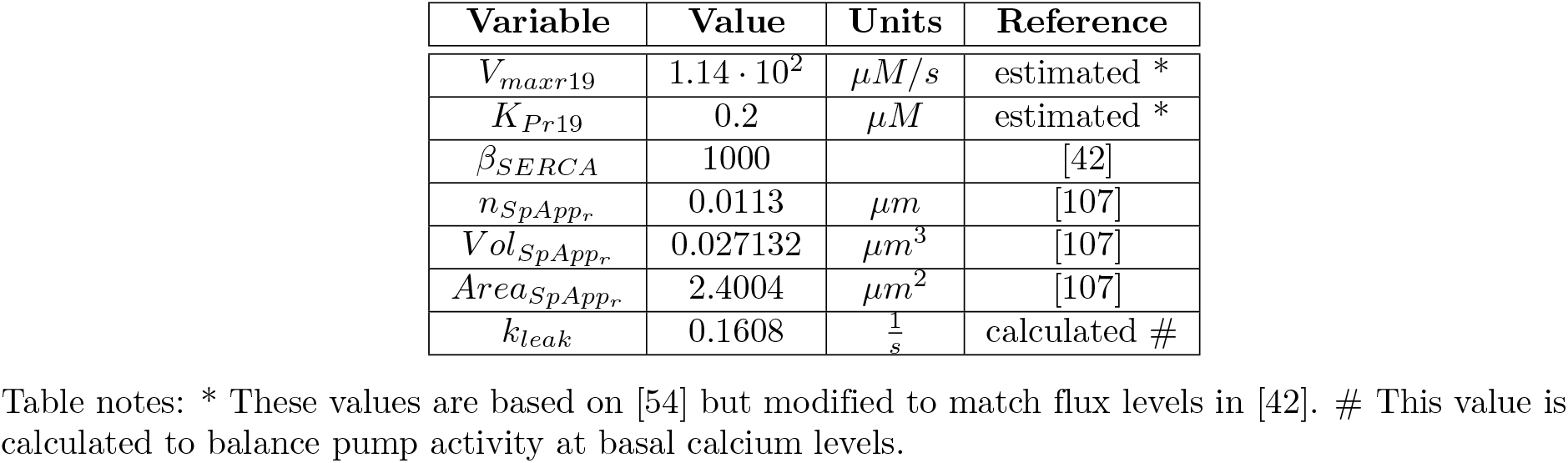
Parameters for SERCA.

| Variable | Value | Units | Reference |
| --- | --- | --- | --- |
| $V_{maxr19}$ | $1.14 \cdot 10^2$ | $\mu M/s$ | estimated * |
| $K_{Pr19}$ | 0.2 | $\mu M$ | estimated * |
| $\beta_{SERCA}$ | 1000 | | [42] |
| $n_{SpAppr}$ | 0.0113 | $\mu m$ | [107] |
| $Vol_{SpAppr}$ | 0.027132 | $\mu m^3$ | [107] |
| $Area_{SpAppr}$ | 2.4004 | $\mu m^2$ | [107] |
| $k_{leak}$ | 0.1608 | $\frac{1}{s}$ | calculated # |

Table notes: \* These values are based on [54] but modified to match flux levels in [42]. # This value is calculated to balance pump activity at basal calcium levels.

#### S1.2.3 Spine Apparatus Membrane

The spine apparatus membrane has SERCA pumps that respond to the increase in cytosolic calcium by intaking cytosolic calcium into the spine apparatus volume. It also has a leak term representing calcium leaking into the cytosol. Therefore, similarly to the plasma membrane, the spine apparatus (denoted as SpApp below for simplicity) has a flux given by

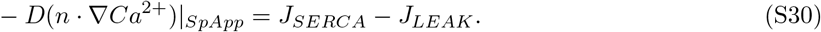

SERCA pump dynamics are based on [54, 118–120] and given by

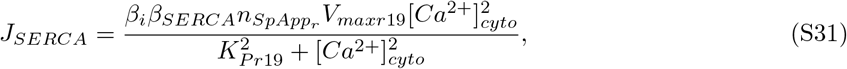

where *β_i_* is defined above in the PM pump section, and *n_SpAppr_* is defined in the same manner as *n*_*PMr*_, but for the spine apparatus volume and membrane surface area. The magnitude of the SERCA pump was based on [42]. The leak term is given by

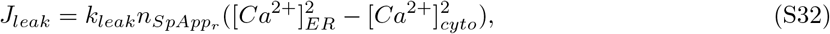

where the leak term is set to balance pump terms at basal calcium concentration [52]. All parameters for the SERCA pumps and leak can be found in Table S7.

## S2 Kinetic parameters used in the model

We have conducted a sensitivity analysis on the numerous parameters in this model for the accumulated calcium (AUC), see Figure S3. We find that the parameters that show significant influence on AUC are well-constrained by experimental observations. The sensitivity analysis was performed in COPASI by converting our spatial model into a compartmental ODE model. Specifically, boundary conditions are converted into volumetric reaction rates through the removal or use of the ‘n’ lengthscale. In addition, fluxes involving the spine apparatus are scaled by the volume ratio 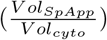 [121]. We also compare our calcium temporal dynamics for our spherical control spine to other existing models of calcium dynamics (Figure S2). Note that while we compare to models with similar stimulus, we do not account for differences in the modeled spine volume. Similarly, we compare our spherical control to experimentally reported calcium transients (Figure 2d). PlotDigitizer was used to trace experimental fluorescence data collected in trials with similar stimulus to our model ([27, 63, 64]). We assumed a linear relationship between fluorescence readout and calcium concentration [65]. However, the explicit relationship between fluorescence readout and calcium concentration is quite complicated and remains an ongoing challenge when comparing fluorescence to concentration. We should also note that upon stimulation of dendritic spines in experimental trials, 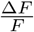 values can range from 0 to 300% percent but depend on the fluorescent calcium probe [27, 122]. We note that some calcium indicators have a weak fluorescence under unstimulated states, so it might appear from some experimental probes that there is no basal calcium concentration. However, use of probes sensitive to unstimulated calcium, such as OGB-1, show the existence of basal levels of calcium [27].

**Fig S2.**
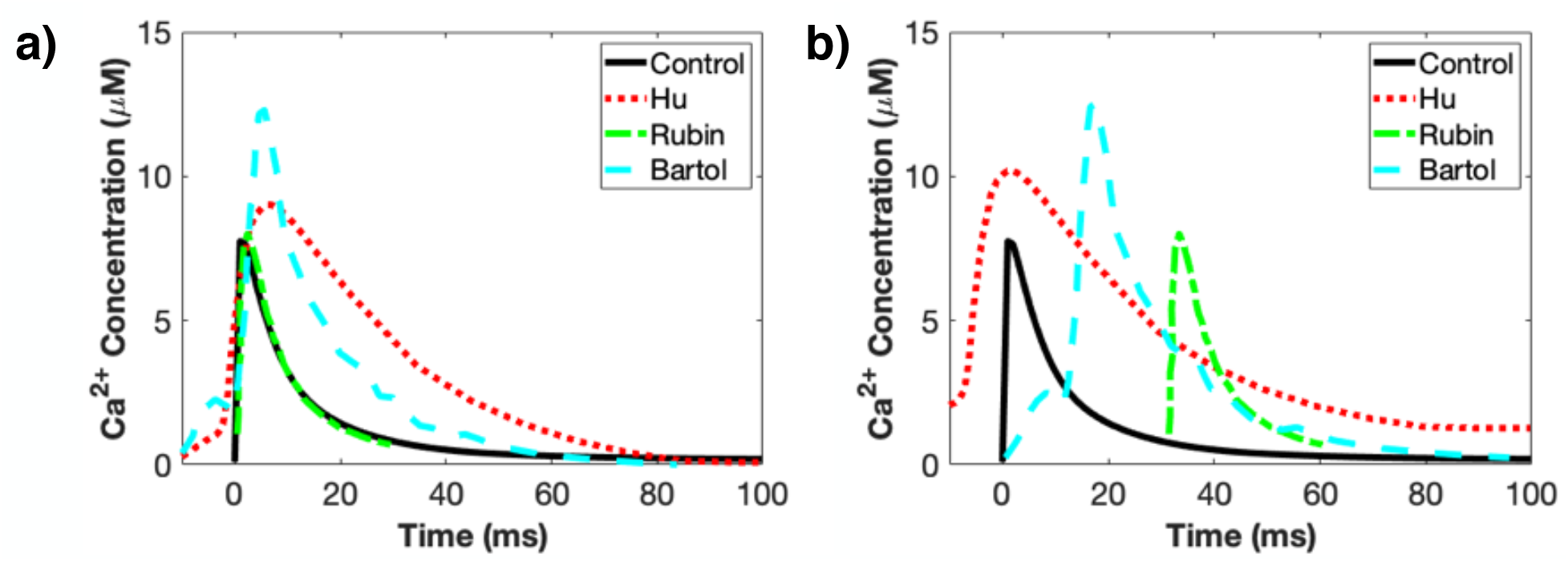
Comparison of calcium temporal dynamics with other existing models. Temporal dynamics of calcium in models with similar stimulus. We plot our spherical control dynamics at the top of the spine versus three existing models Hu et al [42], Rubin et al [123], and Bartol et al [6]. Specifically we are comparing Fig 5 of [42], Fig 1A of [123], and Fig. 7I of [6]. a) We shift all traces to have similar activation starting points and eliminate basal calcium rates at 100ms. We see that our dynamics match Rubin et al very closely. We note that this is a single temporal comparison and we cannot account for differences in spine size across each trace. b) Raw plots as seen in all individual papers.

## S3 Geometric parameters used in the model

For the spherical spine, we use a spine head radius of 237 nm with a spine neck of radius 50 nm. The spherical spine apparatus has a head radius of 80 nm and neck radius of 20 nm. Spine necks were approximately 318 nm long and spine apparatus necks were approximately 363 nm. The geometry used in this model was inspired by [6, 22, 41, 56, 124, 125]. For the ellipsoidal spine and spine apparatus, we set the volumes to be the same as the spherical spine and spherical spine apparatus. Therefore, the ellipsoidal spines had proportionally more spine neck in the final object, with a spine neck of about 354 nm and spine apparatus neck of about 362 nm. For the ellipsoidal shapes, a constant ratio of the a, b, and c axes was kept for both the spine head and spine apparatus. When b is the long axis (length), a is medium axis (width), and c is short axis (height), a ratio of 27:15:7 (b:a:c) was maintained. The volume and surface area measurements of the various geometries used are given in Table S8. Note that the spine volume corresponds to the volume of the cytoplasm. The whole volume of the entire dendritic spine structure would be the combined total of the spine cytoplasmic volume and the spine apparatus volume.

For Figures S4 and S5, we explicitly model a dendritic shaft attached to the spine neck base. We model the dendrite as a cylinder of length 1 *µ*m and radius of 250 nm. We include a basal rate of calcium of 100 nM and have no flux boundary conditions along the entire dendrite. For (Figure S6), we varied the spine neck radius as a cylinder of different radius and height to preserve constant volume. For the smaller neck, the spine neck has a radius of 35 nm and height of 645 nm. For the larger neck, the spine neck has a radius of 60.5 nm and height of 221 nm. We also vary the sizes of the spherical and ellipsoidal spines to elucidate the effect of size on calcium dynamics. The volumes and surface areas of the spherical and ellipsoidal spines can be found in Table S9 and Table S10, respectively. We also vary spine size by varying the spine apparatus size, the volumes and surface areas of the spherical and ellipsoidal spines can be found in Table S11 and Table S12, respectively.

**Fig S3.**
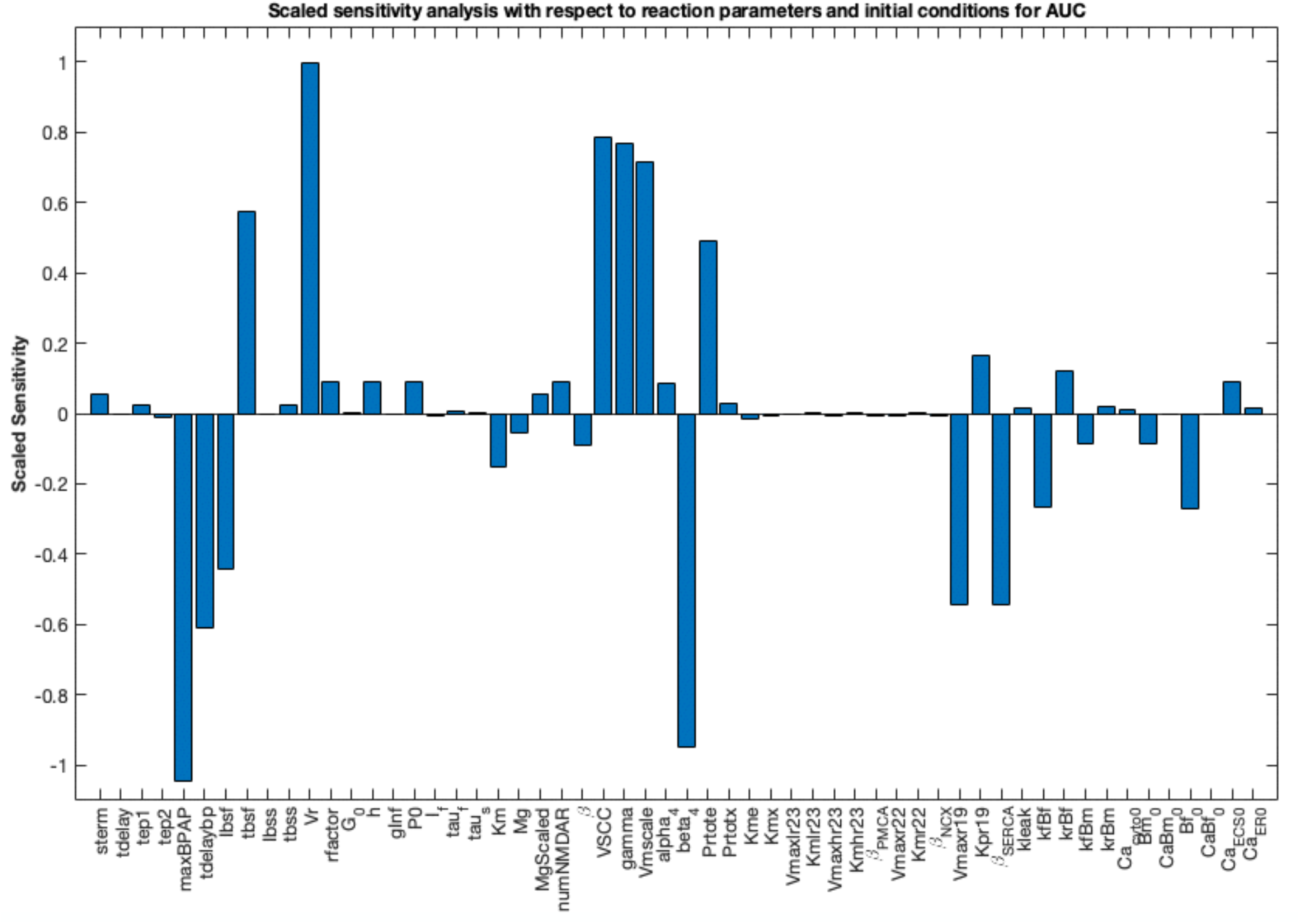
Sensitivity analysis with respect to various initial conditions and reaction parameters. We plot scaled sensitivity for the various reaction parameters and initial conditions in our model converted into a compartmental ordinary differential equations system. Please see Tables S2 to S7 for justification of all parameters, including those that show more sensitivity.

**Table S8.**
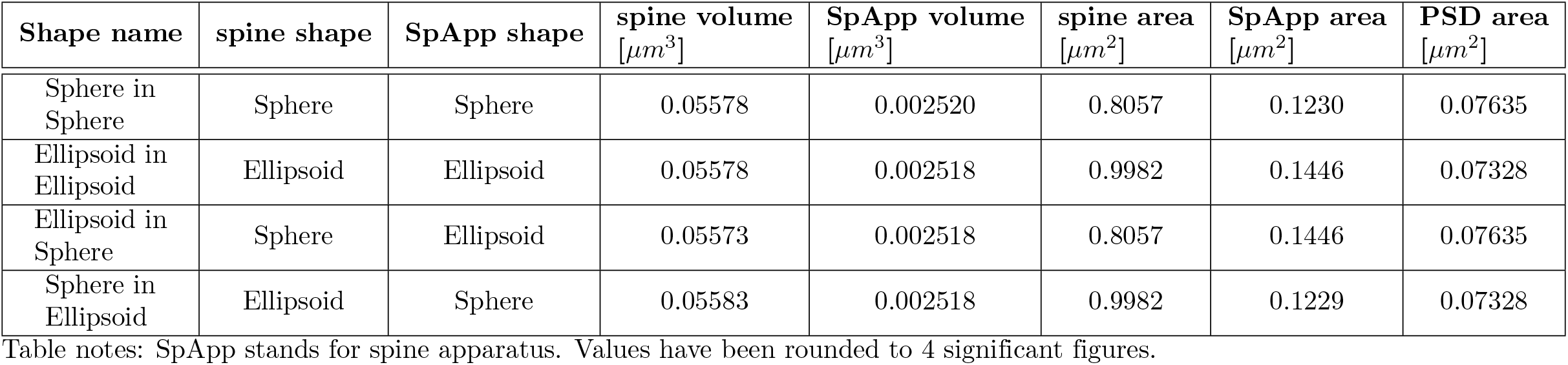
Size and shape parameters of the different geometries

**Table S9.** Size variations for the spherical spine head with spherical spine apparatus.

| Shape and Size | spine volume<br>[$\mu m^3$] | PM area<br>[$\mu m^2$] | SpApp volume<br>[$\mu m^3$] | SpApp area<br>[$\mu m^2$] | PSD area<br>[$\mu m^2$] |
| --- | --- | --- | --- | --- | --- |
| Sphere 50% | 0.02788 | 0.5561 | 0.002520 | 0.1230 | 0.04761 |
| Sphere 75% | 0.04183 | 0.6878 | 0.002520 | 0.1230 | 0.06283 |
| Sphere Control | 0.05578 | 0.8057 | 0.002520 | 0.1230 | 0.07635 |
| Sphere 125% | 0.06973 | 0.9142 | 0.002520 | 0.1230 | 0.08873 |
| Sphere 150% | 0.08367 | 1.016 | 0.002520 | 0.1230 | 0.1003 |

Table notes: SpApp stands for spine apparatus. PM stands for plasma membrane. Values have been rounded to 4 significant figures.

**Table S10.**
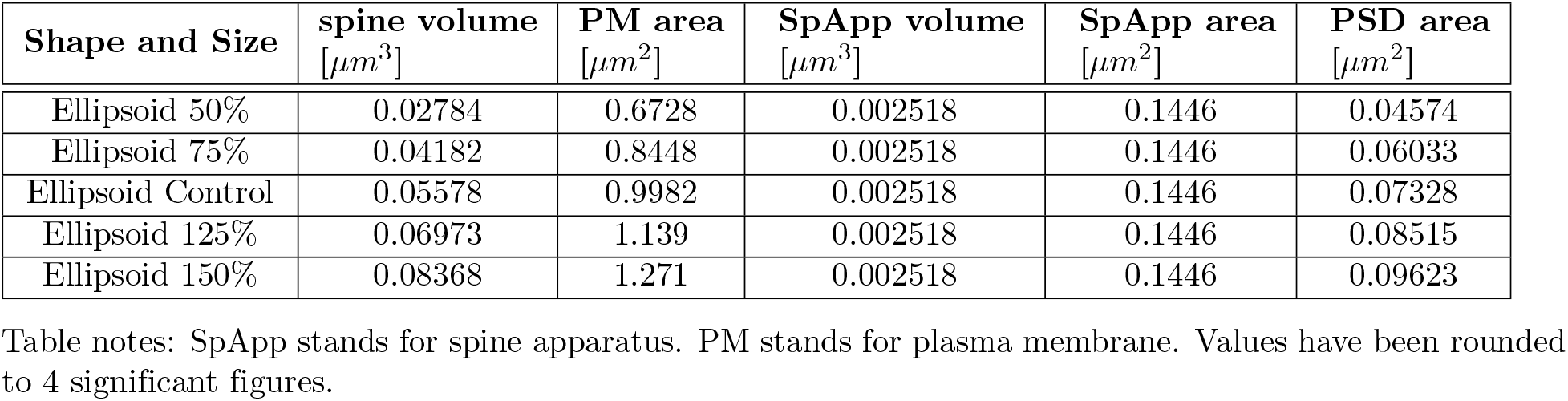
Size variations for the ellipsoidal spine head with ellipsoidal spine apparatus.

| Shape and Size | spine volume<br>[$\mu m^3$] | PM area<br>[$\mu m^2$] | SpApp volume<br>[$\mu m^3$] | SpApp area<br>[$\mu m^2$] | PSD area<br>[$\mu m^2$] |
| --- | --- | --- | --- | --- | --- |
| Ellipsoid 50% | 0.02784 | 0.6728 | 0.002518 | 0.1446 | 0.04574 |
| Ellipsoid 75% | 0.04182 | 0.8448 | 0.002518 | 0.1446 | 0.06033 |
| Ellipsoid Control | 0.05578 | 0.9982 | 0.002518 | 0.1446 | 0.07328 |
| Ellipsoid 125% | 0.06973 | 1.139 | 0.002518 | 0.1446 | 0.08515 |
| Ellipsoid 150% | 0.08368 | 1.271 | 0.002518 | 0.1446 | 0.09623 |

Table notes: SpApp stands for spine apparatus. PM stands for plasma membrane. Values have been rounded to 4 significant figures.

**Table S11.** Size variations for the spherical spine head with spherical spine apparatus when varying spine apparatus volume.

| Shape and Size | spine volume<br>[$\mu m^3$] | PM area<br>[$\mu m^2$] | SpApp volume<br>[$\mu m^3$] | SpApp area<br>[$\mu m^2$] | PSD area<br>[$\mu m^2$] |
| --- | --- | --- | --- | --- | --- |
| Sphere 50% | 0.02784 | 0.8057 | 0.03077 | 0.5004 | 0.07635 |
| Sphere 75% | 0.04183 | 0.8057 | 0.01671 | 0.3447 | 0.07635 |
| Sphere Control | 0.05578 | 0.8057 | 0.002520 | 0.1230 | 0.07635 |

Table notes: SpApp stands for spine apparatus. PM stands for plasma membrane. Values have been rounded to 4 significant figures.

**Table S12.** Size variations for the ellipsoidal spine head with ellipsoidal spine apparatus when varying spine apparatus volume.

| Shape and Size | spine volume<br>[$\mu m^3$] | PM area<br>[$\mu m^2$] | SpApp volume<br>[$\mu m^3$] | SpApp area<br>[$\mu m^2$] | PSD area<br>[$\mu m^2$] |
| --- | --- | --- | --- | --- | --- |
| Ellipsoid 50% | 0.02785 | 0.9982 | 0.03046 | 0.6280 | 0.07328 |
| Ellipsoid 75% | 0.04189 | 0.9982 | 0.01648 | 0.4296 | 0.07328 |
| Ellipsoid Control | 0.05578 | 0.9982 | 0.002518 | 0.1446 | 0.07328 |

Table notes: SpApp stands for spine apparatus. PM stands for plasma membrane. Values have been rounded to 4 significant figures.

## S4 Treatment of the neck base flux can alter calcium dynamics

Thus far, we have treated the dendrite spine as an isolated system. Therefore, we treated the base of the spine neck in the same way as the rest of the plasma membrane, with VSCC causing calcium influx into the spine, while PMCA and NCX pump calcium out of the spine. However, in reality, the dendritic spine is attached to a dendrite. Therefore, we next considered possible fluxes at the base of the spine neck and their effects on calcium dynamics. We consider two other cases to investigate the consequences of different boundary conditions at the base of the spine neck (Figure S4) – i) we clamp the neck base at a concentration of zero micromolar calcium. We see that clamping the base at zero has a dramatic effect on calcium dynamics by forcing a constant spatial gradient (Figure S4a,d) and significantly decreasing AUC and peak concentration (Figure S4b,c). ii) We also explicitly include the dendrite attached to the spine neck. If we explicitly include the spine dendrite, we see that it also creates a spatial gradient, but shows a more sustained calcium transient which leads to a higher AUC compared to the clamped case. We see similar trends for the ellipsoidal spine results (Figure S5).

## S5 Spine neck radius has limited effects on accumulated calcium readout

We have assumed a constant neck radius of 50 nm for all of the simulation thus far. However, the width of the spine neck has been shown as an important determinant of calcium dynamics when comparing larger and smaller spines [43, 77, 87]. Therefore, we varied the spine neck radius (Figure S6) between two extremes allowed by the spine apparatus volume restriction. We found that for our model, the spine neck radius affects the peak calcium and spatial gradients with a higher peak and larger gradient for a thinner neck, but has very limited effects on the accumulated calcium readout. The effect of the spine neck remains of interest however due to its documented ability to affect the diffusion of both cytoplasmic [126] and membrane bound proteins [127–130] and its electrical resistance [131, 132]. See [101] for spine neck variations at a longer timescale and [107] for realistic spine necks.

## S6 Calcium binding proteins are required for physiological temporal dynamics

Calcium dynamics are tightly regulated by binding molecules such that when calcium floods into the spine through NMDAR and VSCC, it is quickly bound by proteins and molecules such as CaM, CaMKII, ATP, and actin [1, 6, 10, 11, 108]. In our model, we consider both fixed and mobile buffers and provide a comparison with exponential decay dynamics to investigate how calcium buffers alter the spatiotemporal dynamics of calcium [10]. Furthermore, the diffusion rate of calcium is of interest due to crowding and cytoskeletonal interactions within the cytoplasm [133]. Therefore, we varied the calcium diffusion coefficient and the mobile buffer concentrations. It is also important to note that during experimental measurements, calcium sensors can bind calcium, attenuating the signaling and giving a lower calcium readout [22, 118]. Therefore, we witness a fairly high calcium concentration since we model the system without these calcium indicators [6].

**Fig S4.**
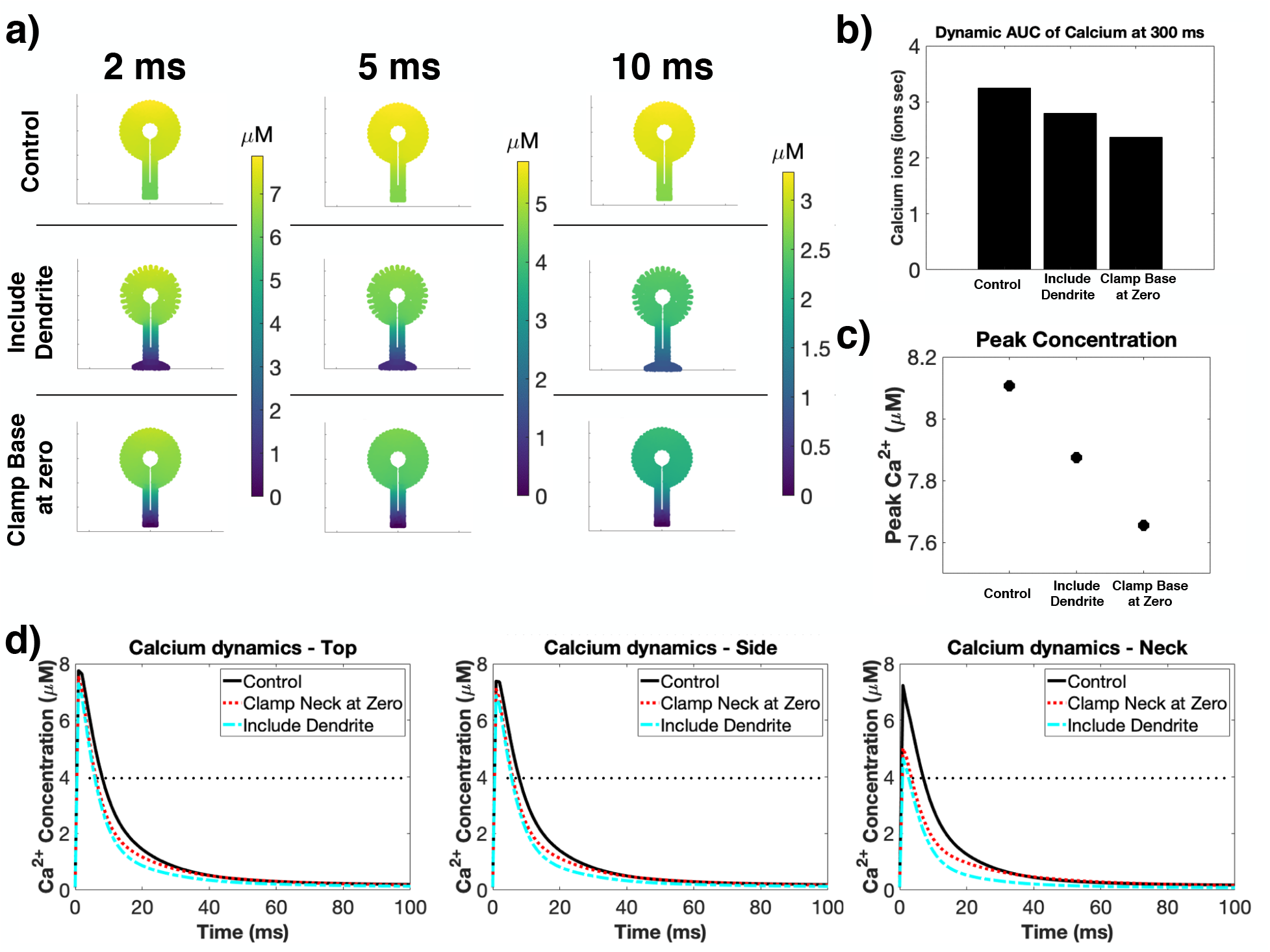
Effect of the boundary conditions at the base of the spine neck on calcium dynamics in spherical spines. a) Spatial dynamics at three different time points (2 ms, 5 ms, and 10 ms) for spherical spines with different boundary conditions at the neck base. AUC (b) and peak concentration (c) for the three conditions shows that the control case has the highest total calcium, then the condition including an explicit dendrite, and finally the base clamped at zero *µM* of calcium has the lowest calcium levels. d) Temporal dynamics at three different locations in the spine (top, side, and neck) show that the boundary conditions at the base of the neck can alter the calcium transients in dendritic spines.

**Fig S5.**
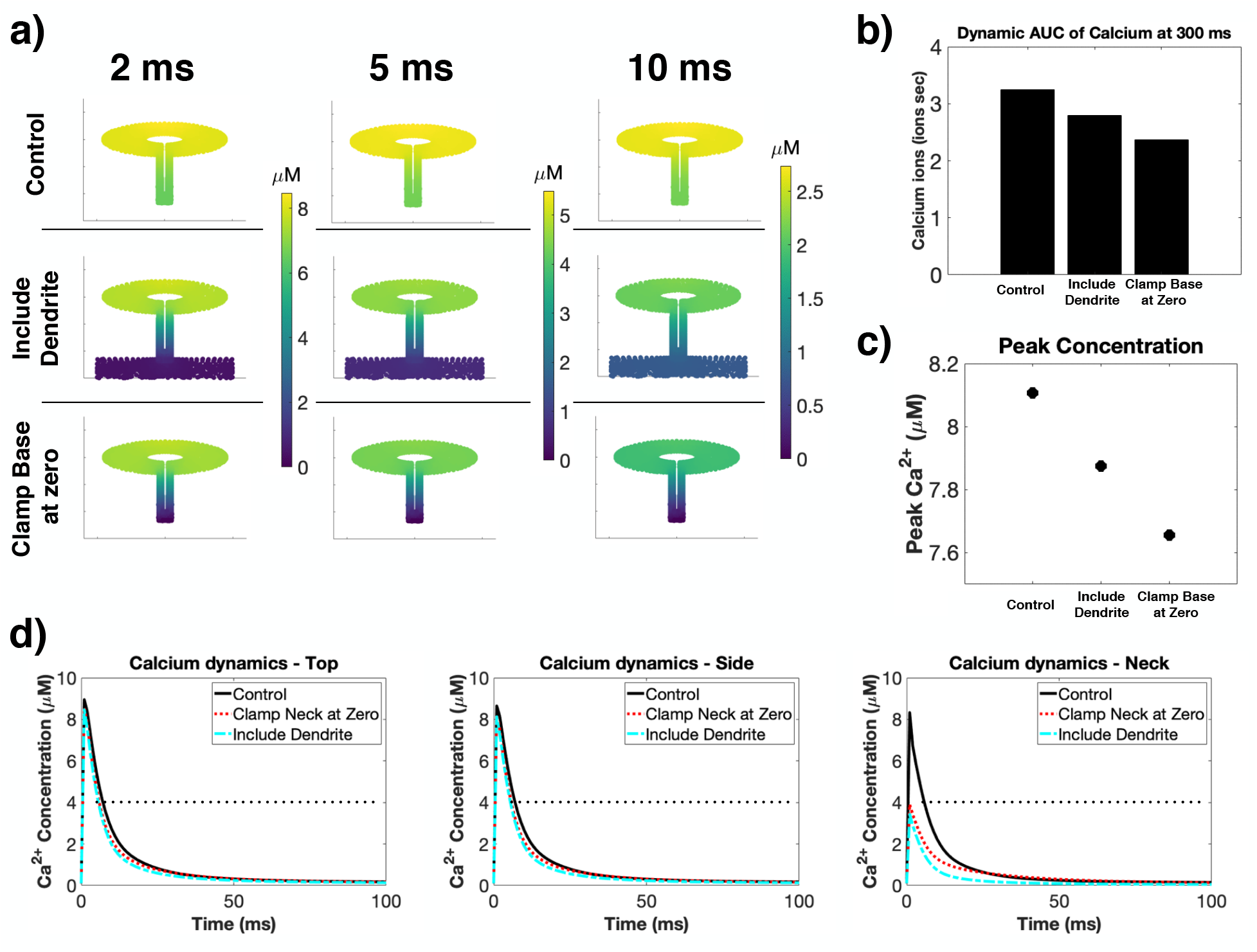
Effect of the boundary conditions at the base of the spine neck on calcium dynamics in ellipsoidal spines. a) Spatial dynamics at three different time points (2 ms, 5 ms, and 10 ms) for ellipsoidal spines with different boundary conditions at the neck base. AUC (b) and peak concentration (c) for the three conditions shows that the control case has the highest total calcium, then the condition including an explicit dendrite, and finally the base clamped at zero *µM* of calcium has the lowest calcium levels, similar to the spherical spine results. d) Temporal dynamics at three different locations in the spine (top, side, and neck) show that the boundary conditions at the base of the neck can alter the calcium transients in dendritic spines at different locations in the spine.

**Fig S6.**
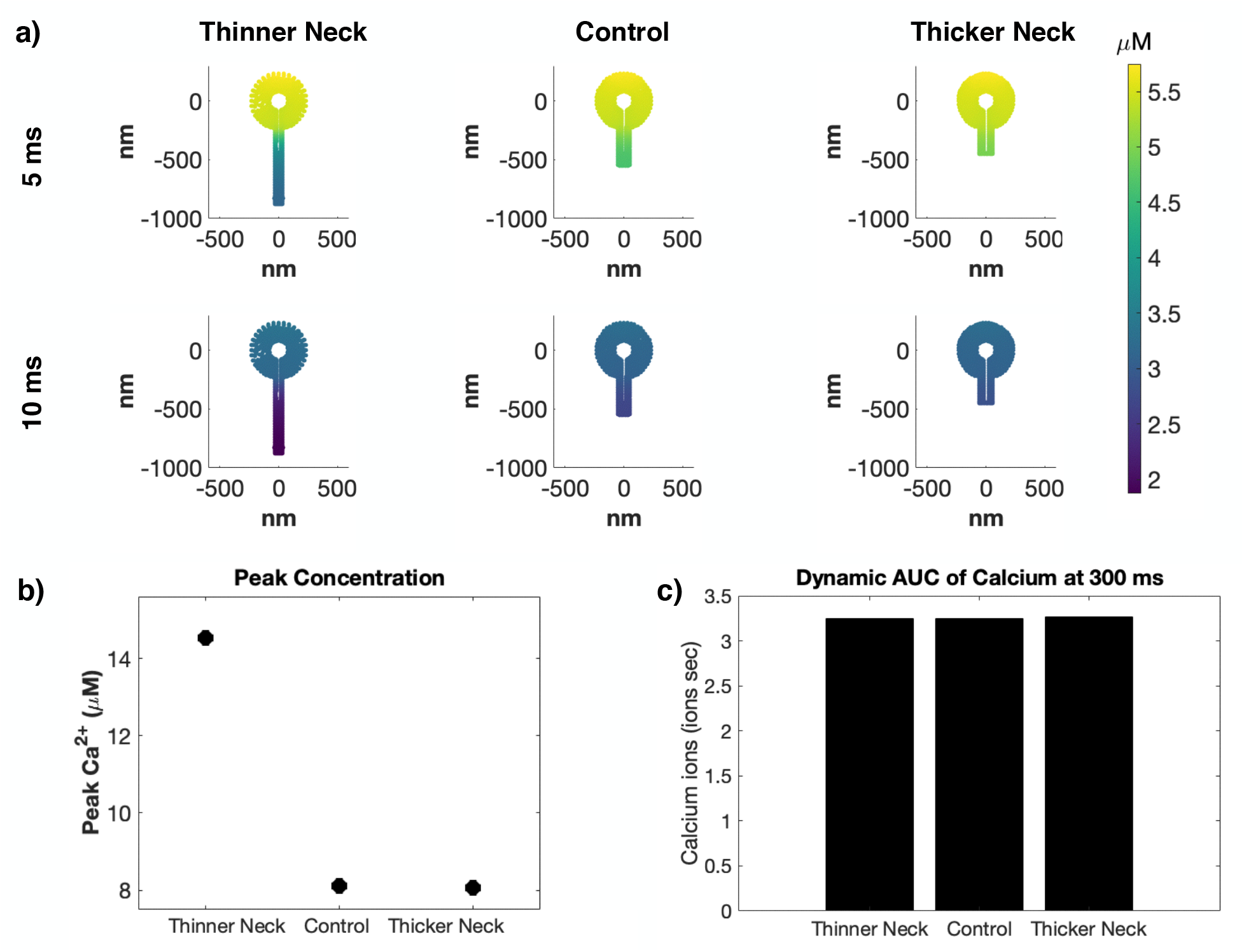
Spine neck radius has limited effects in our isolated spine model. We modeled spines with constant volume but varying neck radius to investigate the effect of neck radius on calcium dynamics. a) Spatial profiles at 5 ms and 10 ms show that spines with longer, thinner necks had a larger concentration gradient than spines with shorter, thicker necks. Spine neck radius had a dramatic effect on peak concentration (b), but had a negligible effect on total calcium for our isolated spine simulations (c).

**Fig S7.**
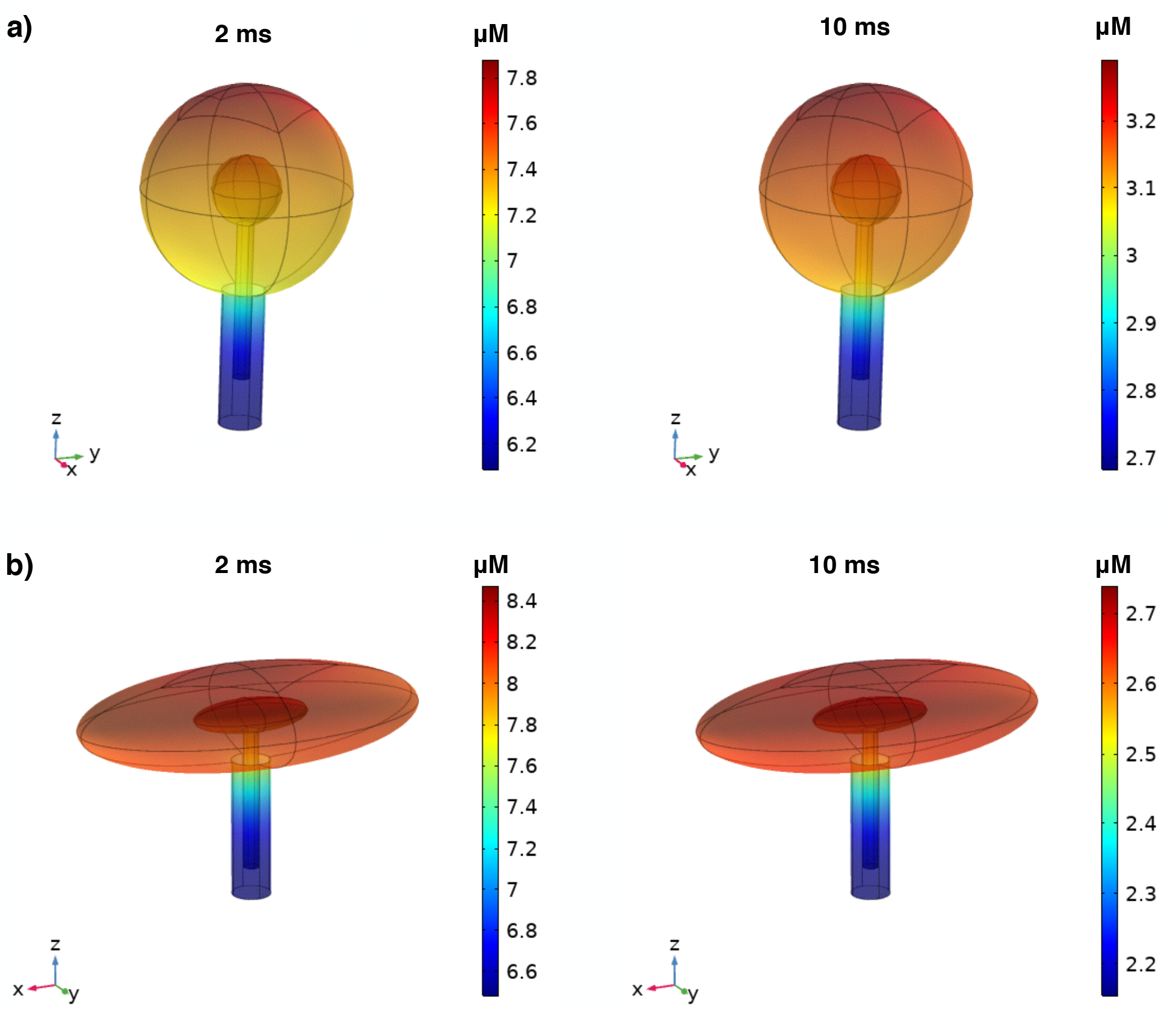
Calcium spatial dynamics in three dimensions. a) Spatial plots of calcium dynamics in the spherical spine at two time points (2 ms, 10 ms). There is a clear spatial gradient from the PSD region at the top of the spine head to the base of the spine neck. b) Spatial plots of calcium dynamics in the ellipsoidal spine at two time points (2 ms, 10 ms). A similar spatial gradient exists for the ellipsoidal spine. The ellipsoidal spine has a larger concentration maximum at 2 ms, while the spherical spine has a larger concentration maximum at 10 ms.

**Fig S8.**
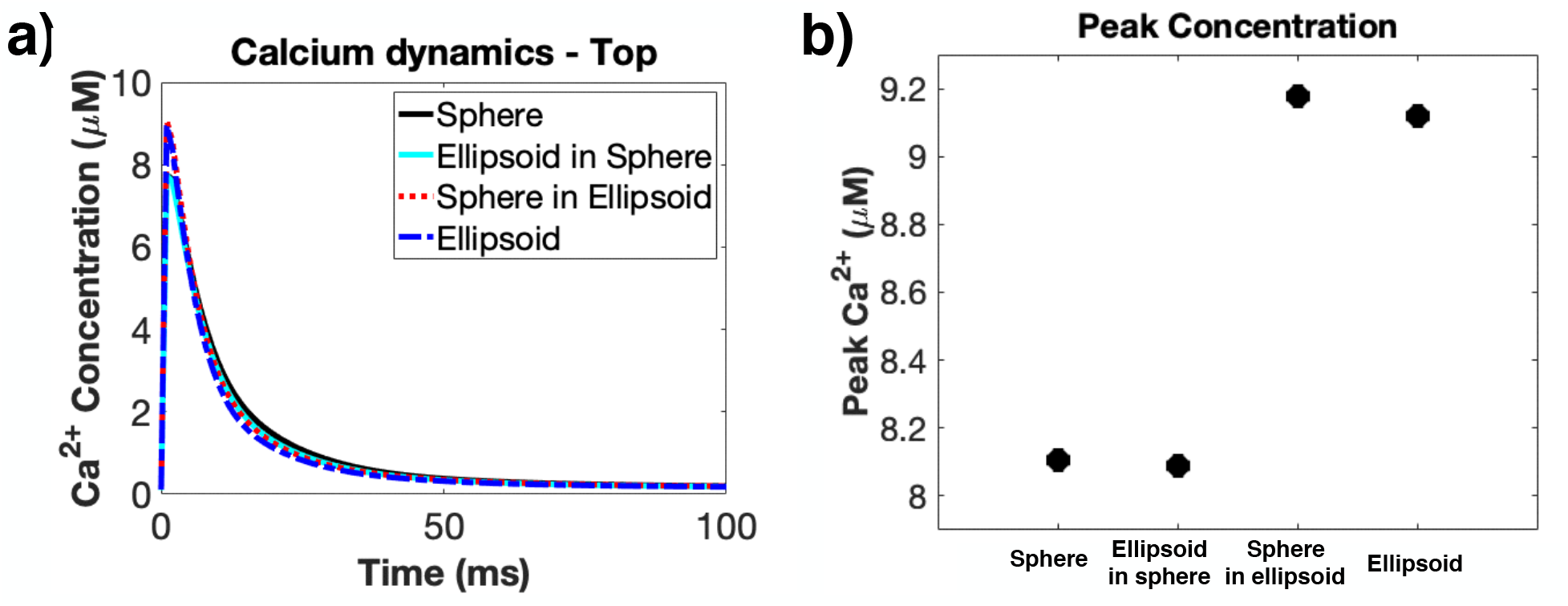
Calcium dynamics in spines of different shapes. a) Calcium temporal dynamics at the top of the spine head for all four shape combinations. All shapes show similar temporal dynamics with a rapid increase in calcium within 2 ms and a decay within 50 ms. b) Peak concentration of calcium for each shape combination. Both spines with ellipsoidal heads show higher max calcium concentration compared to the spines with a spherical head. Spines with an ellipsoidal spine apparatus have a slightly decreased peak compared to the spine with a spherical spine apparatus.

**Fig S9.**
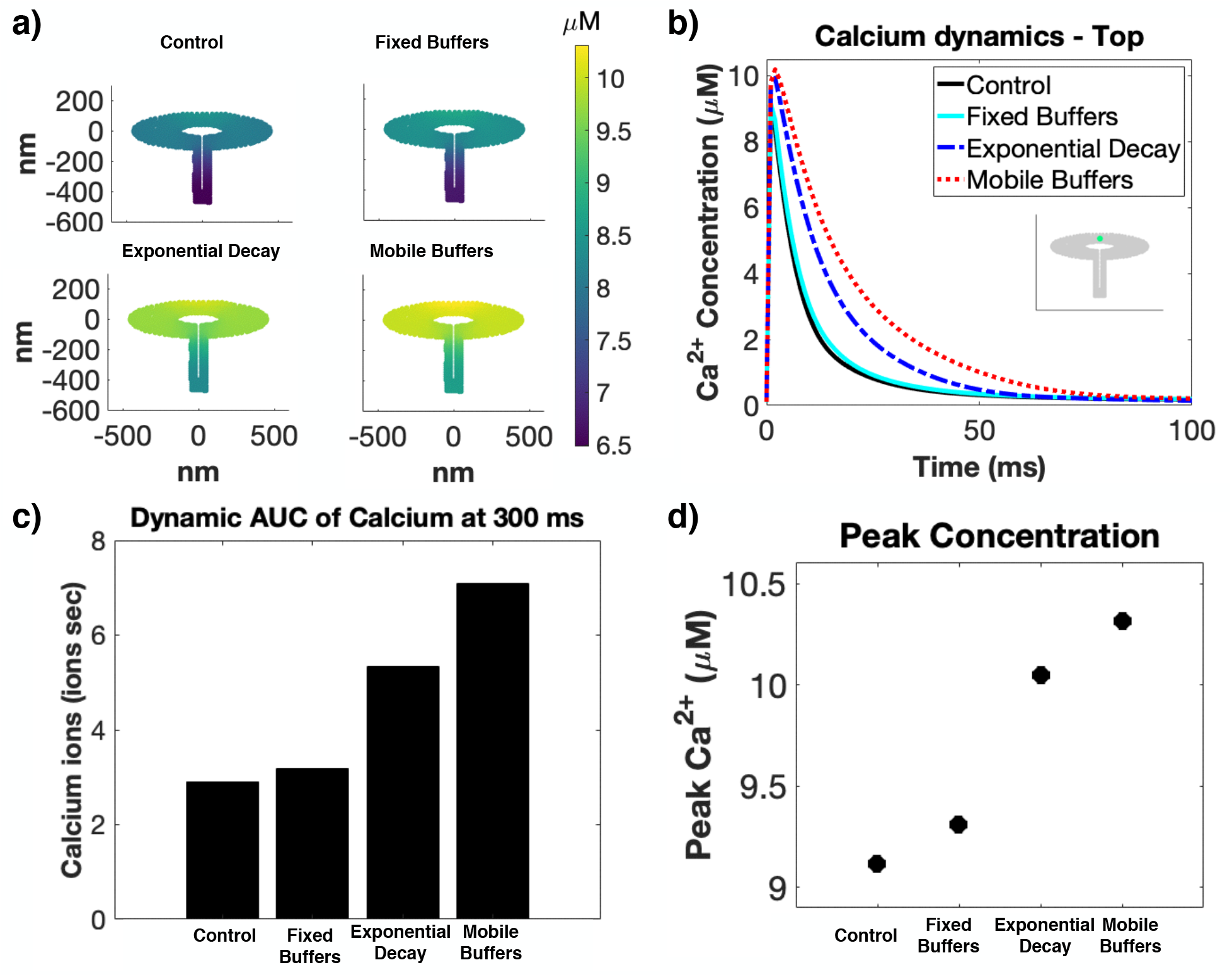
Effect of calcium buffer types on calcium dynamics in ellipsoidal spines. a) Spatial dynamics at 2 ms for ellipsoidal spines with different buffer conditions; control (with both fixed and mobile buffers), only fixed buffers, only mobile buffers, and a lumped exponential decay. While all buffer cases show peak concentrations within about 1.5 *µM* (d), all other quantifications show that they greatly impact the calcium transient decay time (a,b,c). Temporal dynamics (b) show that the control and fixed buffer cases have much faster decay, which translated into lower AUC values (c).

## References

1. Rangamani P, Levy MG, Khan S, Oster G. Paradoxical signaling regulates structural plasticity in dendritic spines. Proceedings of the National Academy of Sciences. 2016;113(36):E5298–E5307.

2. Bourne JN, Harris KM. Balancing structure and function at hippocampal dendritic spines. Annu Rev Neurosci. 2008;31:47–67.

3. Nishiyama J, Yasuda R. Biochemical computation for spine structural plasticity. Neuron. 2015;87(1):63–75.

4. Yasuda R. Biophysics of Biochemical Signaling in Dendritic Spines: Implications in Synaptic Plasticity. Biophysical Journal. 2017;.

5. Denk W, Yuste R, Svoboda K, Tank DW. Imaging calcium dynamics in dendritic spines. Current opinion in neurobiology. 1996;6(3):372–378.

6. Bartol TM, Keller DX, Kinney JP, Bajaj CL, Harris KM, Sejnowski TJ, et al. Computational reconstitution of spine calcium transients from individual proteins. Frontiers in synaptic neuroscience. 2015;7:17.

7. Bloodgood BL, Sabatini BL. Neuronal activity regulates diffusion across the neck of dendritic spines. Science. 2005;310(5749):866–869.

8. Augustine GJ, Santamaria F, Tanaka K. Local calcium signaling in neurons. Neuron. 2003;40(2):331–346.

9. Holmes WR. Is the function of dendritic spines to concentrate calcium? Brain research. 1990;519(1-2):338–342.

10. Yuste R, Majewska A, Holthoff K. From form to function: calcium compartmentalization in dendritic spines. Nature neuroscience. 2000;3(7):653.

11. Segal M. Dendritic spines and long-term plasticity. Nature Reviews Neuroscience. 2005;6(4):277.

12. Jahr CE, Stevens CF. Calcium permeability of the N-methyl-D-aspartate receptor channel in hippocampal neurons in culture. Proceedings of the National Academy of Sciences. 1993;90(24):11573–11577.

13. Shouval HZ, Bear MF, Cooper LN. A unified model of NMDA receptor-dependent bidirectional synaptic plasticity. Proceedings of the National Academy of Sciences. 2002;99(16):10831–10836.

14. Rackham O, Tsaneva-Atanasova K, Ganesh A, Mellor J. A Ca2+-based computational model for NMDA receptor-dependent synaptic plasticity at individual post-synaptic spines in the hippocampus. Frontiers in synaptic neuroscience. 2010;2:31.

15. Jaffe DB, Fisher SA, Brown TH. Confocal laser scanning microscopy reveals voltage-gated calcium signals within hippocampal dendritic spines. Journal of neurobiology. 1994;25(3):220–233.

16. Yuste R, Denk W. Dendritic spines as basic functional units of neuronal integration. Nature. 1995;375(6533):682.

17. Clapham DE. Calcium signaling. Cell. 1995;80(2):259–268.

18. Clapham DE. Calcium signaling. Cell. 2007;131(6):1047–1058.

19. Schmidt H. Three functional facets of calbindin D-28k. Frontiers in molecular neuroscience. 2012;5:25.

20. Holcman D, Schuss Z, Korkotian E. Calcium dynamics in dendritic spines and spine motility. Biophysical journal. 2004;87(1):81–91.

21. Means S, Smith AJ, Shepherd J, Shadid J, Fowler J, Wojcikiewicz RJ, et al. Reaction diffusion modeling of calcium dynamics with realistic ER geometry. Biophysical journal. 2006;91(2):537–557.

22. Higley MJ, Sabatini BL. Calcium signaling in dendritic spines. Cold Spring Harbor perspectives in biology. 2012;4(4):a005686.

23. Loewenstein Y, Sompolinsky H. Temporal integration by calcium dynamics in a model neuron. Nature neuroscience. 2003;6(9):961.

24. Wu Y, Whiteus C, Xu CS, Hayworth KJ, Weinberg RJ, Hess HF, et al. Contacts between the endoplasmic reticulum and other membranes in neurons. Proceedings of the National Academy of Sciences. 2017;114(24):E4859–E4867.

25. Bartol Jr TM, Bromer C, Kinney J, Chirillo MA, Bourne JN, Harris KM, et al. Nanoconnectomic upper bound on the variability of synaptic plasticity. Elife. 2015;4.

26. Kuwajima M, Spacek J, Harris KM. Beyond counts and shapes: studying pathology of dendritic spines in the context of the surrounding neuropil through serial section electron microscopy. Neuroscience. 2013;251:75–89.

27. Sabatini BL, Oertner TG, Svoboda K. The life cycle of Ca2+ ions in dendritic spines. Neuron. 2002;33(3):439–452.

28. Koch C, Zador A. The function of dendritic spines: devices subserving biochemical rather than electrical compartmentalization. Journal of Neuroscience. 1993;13(2):413–422.

29. Hering H, Sheng M. Dentritic spines: structure, dynamics and regulation. Nature Reviews Neuroscience. 2001;2(12):880.

30. Berry KP, Nedivi E. Spine dynamics: are they all the same? Neuron. 2017;96(1):43–55.

31. Jedlicka P, Vlachos A, Schwarzacher SW, Deller T. A role for the spine apparatus in LTP and spatial learning. Behavioural brain research. 2008;192(1):12–19.

32. Basnayake K, Korkotian E, Holcman D. Fast calcium transients in neuronal spines driven by extreme statistics. bioRxiv. 2018;Available from: https://www.biorxiv.org/content/early/2018/04/08/290734.

33. Wuytack F, Raeymaekers L, Missiaen L. Molecular physiology of the SERCA and SPCA pumps. Cell calcium. 2002;32(5-6):279–305.

34. Spacek J, Harris KM. Three-dimensional organization of smooth endoplasmic reticulum in hippocampal CA1 dendrites and dendritic spines of the immature and mature rat. Journal of Neuroscience. 1997;17(1):190–203.

35. Holbro N, Grunditz Å, Oertner TG. Differential distribution of endoplasmic reticulum controls metabotropic signaling and plasticity at hippocampal synapses. Proceedings of the National Academy of Sciences. 2009;106(35):15055–15060.

36. Kotaleski JH, Blackwell KT. Modelling the molecular mechanisms of synaptic plasticity using systems biology approaches. Nature Reviews Neuroscience. 2010;11(4):nrn2807.

37. Gallimore AR, Aricescu AR, Yuzaki M, Calinescu R. A Computational Model for the AMPA Receptor Phosphorylation Master Switch Regulating Cerebellar Long-Term Depression. PLoS Comput Biol. 2016;12(1):e1004664.

38. Byrne MJ, Waxham MN, Kubota Y. The impacts of geometry and binding on CaMKII diffusion and retention in dendritic spines. Journal of computational neuroscience. 2011;31(1):1–12.

39. Wacquier B, Combettes L, Van Nhieu GT, Dupont G. Interplay between intracellular Ca 2+ oscillations and Ca 2+-stimulated mitochondrial metabolism. Scientific reports. 2016;6:19316.

40. Mattson MP, LaFerla FM, Chan SL, Leissring MA, Shepel PN, Geiger JD. Calcium signaling in the ER: its role in neuronal plasticity and neurodegenerative disorders. Trends in neurosciences. 2000;23(5):222–229.

41. Griffith T, Tsaneva-Atanasova K, Mellor JR. Control of Ca2+ influx and calmodulin activation by SK-channels in dendritic spines. PLoS computational biology. 2016;12(5):e1004949.

42. Hu E, Mergenthal A, Bingham CS, Song D, Bouteiller JM, Berger TW. A glutamatergic spine model to enable multi-scale modeling of nonlinear calcium dynamics. Frontiers in Computational Neuroscience. 2018;12.

43. Arellano JI, Benavides-Piccione R, DeFelipe J, Yuste R. Ultrastructure of dendritic spines: correlation between synaptic and spine morphologies. Frontiers in neuroscience. 2007;1:10.

44. Ostroff LE, Botsford B, Gindina S, Cowansage KK, LeDoux JE, Klann E, et al. Accumulation of polyribosomes in dendritic spine heads, but not bases and necks, during memory consolidation depends on cap-dependent translation initiation. Journal of Neuroscience. 2017;37(7):1862–1872.

45. Segal M, Vlachos A, Korkotian E. The spine apparatus, synaptopodin, and dendritic spine plasticity. The Neuroscientist. 2010;16(2):125–131.

46. Lee SJR, Escobedo-Lozoya Y, Szatmari EM, Yasuda R. Activation of CaMKII in single dendritic spines during long-term potentiation. Nature. 2009;458(7236):299.

47. Sabatini BL, Maravall M, Svoboda K. Ca2+ signaling in dendritic spines. Current opinion in neurobiology. 2001;11(3):349–356.

48. Schmidt H, Eilers J. Spine neck geometry determines spino-dendritic cross-talk in the presence of mobile endogenous calcium binding proteins. Journal of computational neuroscience. 2009;27(2):229–243.

49. Matthews EA, Dietrich D. Buffer mobility and the regulation of neuronal calcium domains. Frontiers in cellular neuroscience. 2015;9:48.

50. Fifkova E. A possible mechanism of morphometric changes in dendritic spines induced by stimulation. Cellular and molecular neurobiology. 1985;5(1-2):47–63.

51. Fifková E, Markham JA, Delay RJ. Calcium in the spine apparatus of dendritic spines in the dentate molecular layer. Brain research. 1983;266(1):163–168.

52. Futagi D, Kitano K. Ryanodine-receptor-driven intracellular calcium dynamics underlying spatial association of synaptic plasticity. Journal of computational neuroscience. 2015;39(3):329–347.

53. Maurya MR, Subramaniam S. A kinetic model for calcium dynamics in RAW 264.7 cells: 1. Mechanisms, parameters, and subpopulational variability. Biophysical journal. 2007;93(3):709–728.

54. Calizo RC, Ron A, Hu M, Bhattacya S, Janssen WGM, Hone J, et al. Curvature regulates subcellular organelle location to control intracellular signal propagation. bioRxiv. 2017;Available from: https://www.biorxiv.org/content/early/2017/07/11/161950.

55. Cugno A, Bartol TM, Sejnowski TJ, Iyengar R, Rangamani P. Geometric principles of second messenger dynamics in dendritic spines. bioRxiv. 2018;Available from: https://www.biorxiv.org/content/early/2018/10/20/444489.

56. Paulin JJ, Haslehurst P, Fellows AD, Liu W, Jackson JD, Joel Z, et al. Large and small dendritic spines serve different interacting functions in hippocampal synaptic plasticity and homeostasis. Neural plasticity. 2016;2016.

57. Ngo-Anh TJ, Bloodgood BL, Lin M, Sabatini BL, Maylie J, Adelman JP. SK channels and NMDA receptors form a Ca 2+-mediated feedback loop in dendritic spines. Nature neuroscience. 2005;8(5):642.

58. Matthews EA, Schoch S, Dietrich D. Tuning local calcium availability: cell-type-specific immobile calcium buffer capacity in hippocampal neurons. Journal of Neuroscience. 2013;33(36):14431–14445.

59. Basak R, Narayanan R. Active dendrites regulate the spatiotemporal spread of signaling microdomains. PLoS computational biology. 2018;14(11):e1006485.

60. Heinrich R, Neel BG, Rapoport TA. Mathematical models of protein kinase signal transduction. Molecular cell. 2002;9(5):957–970.

61. Atay O, Skotheim JM. Spatial and temporal signal processing and decision making by MAPK pathways. J Cell Biol. 2017;216(2):317–330.

62. Gorman RP, Sejnowski TJ. Analysis of hidden units in a layered network trained to classify sonar targets. Neural networks. 1988;1(1):75–89.

63. Segal M, Korkotian E. Endoplasmic reticulum calcium stores in dendritic spines. Frontiers in neuroanatomy. 2014;8:64.

64. Hoogland TM, Saggau P. Facilitation of L-type Ca2+ channels in dendritic spines by activation of *β*2 adrenergic receptors. Journal of Neuroscience. 2004;24(39):8416–8427.

65. Yasuda R, Nimchinsky EA, Scheuss V, Pologruto TA, Oertner TG, Sabatini BL, et al. Imaging calcium concentration dynamics in small neuronal compartments. Sci STKE. 2004;2004(219):pl5–pl5.

66. Kasai H, Fukuda M, Watanabe S, Hayashi-Takagi A, Noguchi J. Structural dynamics of dendritic spines in memory and cognition. Trends in neurosciences. 2010;33(3):121–129.

67. Knott GW, Holtmaat A, Wilbrecht L, Welker E, Svoboda K. Spine growth precedes synapse formation in the adult neocortex in vivo. Nature neuroscience. 2006;9(9):1117.

68. Nixon-Abell J, Obara CJ, Weigel AV, Li D, Legant WR, Xu CS, et al. Increased spatiotemporal resolution reveals highly dynamic dense tubular matrices in the peripheral ER. Science. 2016;354(6311):aaf3928.

69. Deller T, Bas Orth C, Vlachos A, Merten T, Del Turco D, Dehn D, et al. Plasticity of synaptopodin and the spine apparatus organelle in the rat fascia dentata following entorhinal cortex lesion. Journal of Comparative Neurology. 2006;499(3):471–484.

70. Wilson C, Groves P, Kitai S, Linder J. Three-dimensional structure of dendritic spines in the rat neostriatum. Journal of Neuroscience. 1983;3(2):383–388.

71. Sabatini BL, Svoboda K. Analysis of calcium channels in single spines using optical fluctuation analysis. Nature. 2000;408(6812):589.

72. Siesjö BK. Calcium in the brain under physiological and pathological conditions. European neurology. 1990;30(Suppl. 2):3–9.

73. Keener JP, Sneyd J. Mathematical physiology. vol. 1. Springer; 1998.

74. Yuste R. Dendritic spines. MIT press; 2010.

75. Rochefort NL, Konnerth A. Dendritic spines: from structure to in vivo function. EMBO reports. 2012;13(8):699–708.

76. Murakoshi H, Yasuda R. Postsynaptic signaling during plasticity of dendritic spines. Trends in neurosciences. 2012;35(2):135–143.

77. Noguchi J, Matsuzaki M, Ellis-Davies GC, Kasai H. Spine-neck geometry determines NMDA receptor-dependent Ca2+ signaling in dendrites. Neuron. 2005;46(4):609–622.

78. Matsuzaki M, Ellis-Davies GC, Nemoto T, Miyashita Y, Iino M, Kasai H. Dendritic spine geometry is critical for AMPA receptor expression in hippocampal CA1 pyramidal neurons. Nature neuroscience. 2001;4(11):1086.

79. Lee KF, Soares C, Béïque JC. Examining form and function of dendritic spines. Neural plasticity. 2012;2012.

80. Béïque JC, Na Y, Kuhl D, Worley PF, Huganir RL. Arc-dependent synapse-specific homeostatic plasticity. Proceedings of the National Academy of Sciences. 2011;108(2):816–821.

81. Turrigiano G. Too many cooks? Intrinsic and synaptic homeostatic mechanisms in cortical circuit refinement. Annual review of neuroscience. 2011;34:89–103.

82. Lee MC, Yasuda R, Ehlers MD. Metaplasticity at single glutamatergic synapses. Neuron. 2010;66(6):859–870.

83. Zhou Q, Sheng M. NMDA receptors in nervous system diseases. Neuropharmacology. 2013;74:69–75.

84. Hotulainen P, Hoogenraad CC. Actin in dendritic spines: connecting dynamics to function. The Journal of cell biology. 2010;189(4):619–629.

85. Santamaria F, Wils S, De Schutter E, Augustine GJ. Anomalous diffusion in Purkinje cell dendrites caused by spines. Neuron. 2006;52(4):635–648.

86. Ouyang Y, Wong M, Capani F, Rensing N, Lee CS, Liu Q, et al. Transient decrease in F-actin may be necessary for translocation of proteins into dendritic spines. European Journal of Neuroscience. 2005;22(12):2995–3005.

87. Araya R, Vogels TP, Yuste R. Activity-dependent dendritic spine neck changes are correlated with synaptic strength. Proceedings of the National Academy of Sciences. 2014;p. 201321869.

88. Voorsluijs V, Dawson SP, De Decker Y, Dupont G. Deterministic limit of intracellular calcium spikes. Physical review letters. 2019;122(8):088101.

89. Yeung LC, Shouval HZ, Blais BS, Cooper LN. Synaptic homeostasis and input selectivity follow from a calcium-dependent plasticity model. Proceedings of the National Academy of Sciences. 2004;101(41):14943–14948.

90. Cummings JA, Mulkey RM, Nicoll RA, Malenka RC. Ca2+ signaling requirements for long-term depression in the hippocampus. Neuron. 1996;16(4):825–833.

91. Malenka RC, Kauer JA, Zucker RS, Nicoll RA. Postsynaptic calcium is sufficient for potentiation of hippocampal synaptic transmission. Science. 1988;242(4875):81–84.

92. Song S, Miller KD, Abbott LF. Competitive Hebbian learning through spike-timing-dependent synaptic plasticity. Nature neuroscience. 2000;3(9):919.

93. Izhikevich EM. Solving the distal reward problem through linkage of STDP and dopamine signaling. Cerebral cortex. 2007;17(10):2443–2452.

94. Bush D, Jin Y. Calcium control of triphasic hippocampal STDP. Journal of computational neuroscience. 2012;33(3):495–514.

95. Sejnowski T. Statistical constraints on synaptic plasticity. Journal of theoretical biology. 1977;69(2):385–389.

96. Standage D, Trappenberg T, Blohm G. Calcium-dependent calcium decay explains STDP in a dynamic model of hippocampal synapses. PloS one. 2014;9(1):e86248.

97. Cruz-Albrecht JM, Yung MW, Srinivasa N. Energy-efficient neuron, synapse and STDP integrated circuits. IEEE transactions on biomedical circuits and systems. 2012;6(3):246–256.

98. Yasuda H, Barth AL, Stellwagen D, Malenka RC. A developmental switch in the signaling cascades for LTP induction. Nature neuroscience. 2003;6(1):15.

99. Oertner TG, Matus A. Calcium regulation of actin dynamics in dendritic spines. Cell calcium. 2005;37(5):477–482.

100. Miermans C, Kusters R, Hoogenraad C, Storm C. Biophysical model of the role of actin remodeling on dendritic spine morphology. PloS one. 2017;12(2):e0170113.

101. Ohadi D, Rangamani P. Geometric control of frequency modulation of cAMP oscillations due to Ca2+-bursts in dendritic spines. bioRxiv. 2019;Available from: https://www.biorxiv.org/content/early/2019/01/15/520643.

102. Ohadi D, Schmitt DL, Calabrese B, Halpain S, Zhang J, Rangamani P. Computational modeling reveals frequency modulation of calcium-cAMP/PKA pathway in dendritic spines. bioRxiv. 2019;Available from: https://www.biorxiv.org/content/early/2019/01/16/521740.

103. Majewska A, Tashiro A, Yuste R. Regulation of spine calcium dynamics by rapid spine motility. Journal of Neuroscience. 2000;20(22):8262–8268.

104. Schmidt H, Kunerth S, Wilms C, Strotmann R, Eilers J. Spino-dendritic cross-talk in rodent Purkinje neurons mediated by endogenous Ca2+-binding proteins. The Journal of physiology. 2007;581(2):619–629.

105. Herz AV, Gollisch T, Machens CK, Jaeger D. Modeling single-neuron dynamics and computations: a balance of detail and abstraction. science. 2006;314(5796):80–85.

106. Rangamani P, Lipshtat A, Azeloglu EU, Calizo RC, Hu M, Ghassemi S, et al. Decoding information in cell shape. Cell. 2013;154(6):1356–1369.

107. Lee CT, Laughlin JG, Angliviel de La Beaumelle N, Amaro R, McCammon JA, Ramamoorthi R, et al. GAMer 2: A System for 3D Mesh Processing of Cellular Electron Micrographs. bioRxiv. 2019;Available from: https://www.biorxiv.org/content/early/2019/01/29/534479.

108. Ivings L, Pennington SR, Jenkins R, Weiss JL, Burgoyne RD. Identification of Ca2+-dependent binding partners for the neuronal calcium sensor protein neurocalcin *δ*: interaction with actin, clathrin and tubulin. Biochemical Journal. 2002;363(3):599–608.

109. Cornelisse LN, van Elburg RA, Meredith RM, Yuste R, Mansvelder HD. High speed two-photon imaging of calcium dynamics in dendritic spines: consequences for spine calcium kinetics and buffer capacity. PloS one. 2007;2(10):e1073.

110. Robinson R, Stokes R. Electrolyte Solutions, Butterworths Scientific Publications. London; 1959.

111. Naraghi M, Neher E. Linearized buffered Ca2+ diffusion in microdomains and its implications for calculation of [Ca2+] at the mouth of a calcium channel. Journal of Neuroscience. 1997;17(18):6961–6973.

112. Schwaller B. Cytosolic Ca2+ buffers. Cold Spring Harbor perspectives in biology. 2010;p. a004051.

113. Nimchinsky EA, Yasuda R, Oertner TG, Svoboda K. The number of glutamate receptors opened by synaptic stimulation in single hippocampal spines. Journal of Neuroscience. 2004;24(8):2054–2064.

114. Reuveni I, Ghosh S, Barkai E. Real Time Multiplicative Memory Amplification Mediated by Whole-Cell Scaling of Synaptic Response in Key Neurons. PLoS computational biology. 2017;13(1):e1005306.

115. Fuenzalida M, de Sevilla DF, Buño W. Changes of the EPSP waveform regulate the temporal window for spike-timing-dependent plasticity. Journal of Neuroscience. 2007;27(44):11940–11948.

116. Froemke RC, Tsay IA, Raad M, Long JD, Dan Y. Contribution of individual spikes in burst-induced long-term synaptic modification. Journal of neurophysiology. 2006;.

117. Breit M, Kessler M, Stepniewski M, Vlachos A, Queisser G. Spine-to-Dendrite Calcium Modeling Discloses Relevance for Precise Positioning of Ryanodine Receptor-Containing Spine Endoplasmic Reticulum. Scientific reports. 2018;8(1):15624.

118. Fink CC, Slepchenko B, Moraru II, Watras J, Schaff JC, Loew LM. An image-based model of calcium waves in differentiated neuroblastoma cells. Biophysical Journal. 2000;79(1):163–183.

119. Johenning FW, Theis AK, Pannasch U, Rückl M, Rüdiger S, Schmitz D. Ryanodine receptor activation induces long-term plasticity of spine calcium dynamics. PLoS biology. 2015;13(6):e1002181.

120. Sneyd J, Tsaneva-Atanasova K, Bruce J, Straub S, Giovannucci D, Yule D. A model of calcium waves in pancreatic and parotid acinar cells. Biophysical journal. 2003;85(3):1392–1405.

121. Dupont G, Falcke M, Kirk V, Sneyd J. Models of calcium signalling. vol. 43. Springer; 2016.

122. Rietdorf K, Chehab T, Allman S, Bootman MD. Novel improved Ca 2+ indicator dyes on the market-a comparative study of novel Ca 2+ indicators with fluo-4. 2014;.

123. Rubin JE, Gerkin RC, Bi GQ, Chow CC. Calcium time course as a signal for spike-timing–dependent plasticity. Journal of neurophysiology. 2005;93(5):2600–2613.

124. Attardo A, Fitzgerald JE, Schnitzer MJ. Impermanence of dendritic spines in live adult CA1 hippocampus. Nature. 2015;523(7562):592.

125. Kusters R, Kapitein LC, Hoogenraad CC, Storm C. Shape-induced asymmetric diffusion in dendritic spines allows efficient synaptic AMPA receptor trapping. Biophysical journal. 2013;105(12):2743–2750.

126. Adrian M, Kusters R, Wierenga CJ, Storm C, Hoogenraad CC, Kapitein LC. Barriers in the brain: resolving dendritic spine morphology and compartmentalization. Frontiers in neuroanatomy. 2014;8:142.

127. Ashby MC, Maier SR, Nishimune A, Henley JM. Lateral diffusion drives constitutive exchange of AMPA receptors at dendritic spines and is regulated by spine morphology. Journal of Neuroscience. 2006;26(26):7046–7055.

128. Yang G, Huang A, Zhu S, Xiong W. It is time to move: role of lateral diffusion in AMPA receptor trafficking. Journal of Neuroscience. 2006;26(36):9082–9083.

129. Ewers H, Tada T, Petersen JD, Racz B, Sheng M, Choquet D. A septin-dependent diffusion barrier at dendritic spine necks. PloS one. 2014;9(12):e113916.

130. Adrian M, Kusters R, Storm C, Hoogenraad CC, Kapitein LC. Probing the Interplay between Dendritic Spine Morphology and Membrane-Bound Diffusion. Biophysical Journal. 2017;.

131. Segev I, Rall W. Excitable dendrites and spines: earlier theoretical insights elucidate recent direct observations. Trends in neurosciences. 1998;21(11):453–460.

132. Yuste R. Electrical compartmentalization in dendritic spines. Annual review of neuroscience. 2013;36:429–449.

133. Biess A, Korkotian E, Holcman D. Barriers to diffusion in dendrites and estimation of calcium spread following synaptic inputs. PLoS computational biology. 2011;7(10):e1002182.

